# Monocarboxylate transporter 2 is required for the maintenance of myelin and axonal integrity by oligodendrocytes

**DOI:** 10.1101/2025.01.10.632306

**Authors:** Leire Izagirre-Urizar, Luna Mora-Huerta, Irene Soler-Saez, Raquel Morales-Gallel, Maria-Jose Ulloa-Navas, Juan-Carlos Chara, Stefano Calovi, Cyrille Deboux, Laura Merino-Cacho, Citlalli Netzahualcoyotzi, Maria Domercq, José L. Zugaza, Luc Pellerin, Francisco Garcia-Garcia, Jose-Manuel Garcia-Verdugo, Carlos Matute, Brahim Nait-Oumesmar, Vanja Tepavcevic

## Abstract

Neurodegenerative pathologies including multiple sclerosis (MS) are consistently associated with energy deficit in the central nervous system (CNS). This might directly impact myelinating oligodendrocytes as these are particularly vulnerable to metabolic insults. Importantly, oligodendroglial dysfunction and myelin alterations occur in most, if not all neurodegenerative diseases, and are associated with axonal pathology/loss. Thus, elucidating metabolic mechanisms required for oligodendroglial myelin maintenance and axonal support might be crucial to identify therapeutic targets to achieve neuroprotection. While monocarboxylates are important energy fuels for the CNS, their role in myelinating oligodendrocyte function remains unclear. Here we show that, just like neurons, myelinating oligodendrocytes express high affinity monocarboxylate transporter 2 (MCT2) both in mice and humans, which is downregulated in progressive MS. While deletion of MCT2 in mouse oligodendrocytes did not affect the survival of these cells, it resulted in downregulation of lipid synthesis-associated enzymes and failure of myelin maintenance. Moreover, axonal upregulation of lactate dehydrogenase A concomitant with axonal damage was observed but could be alleviated by ketogenic diet. We conclude that oligodendroglial MCT2 is required for myelin maintenance and axonal support, which becomes altered in progressive MS, but may be compensated for by specific metabolic therapies.

**Graphical abstract:** 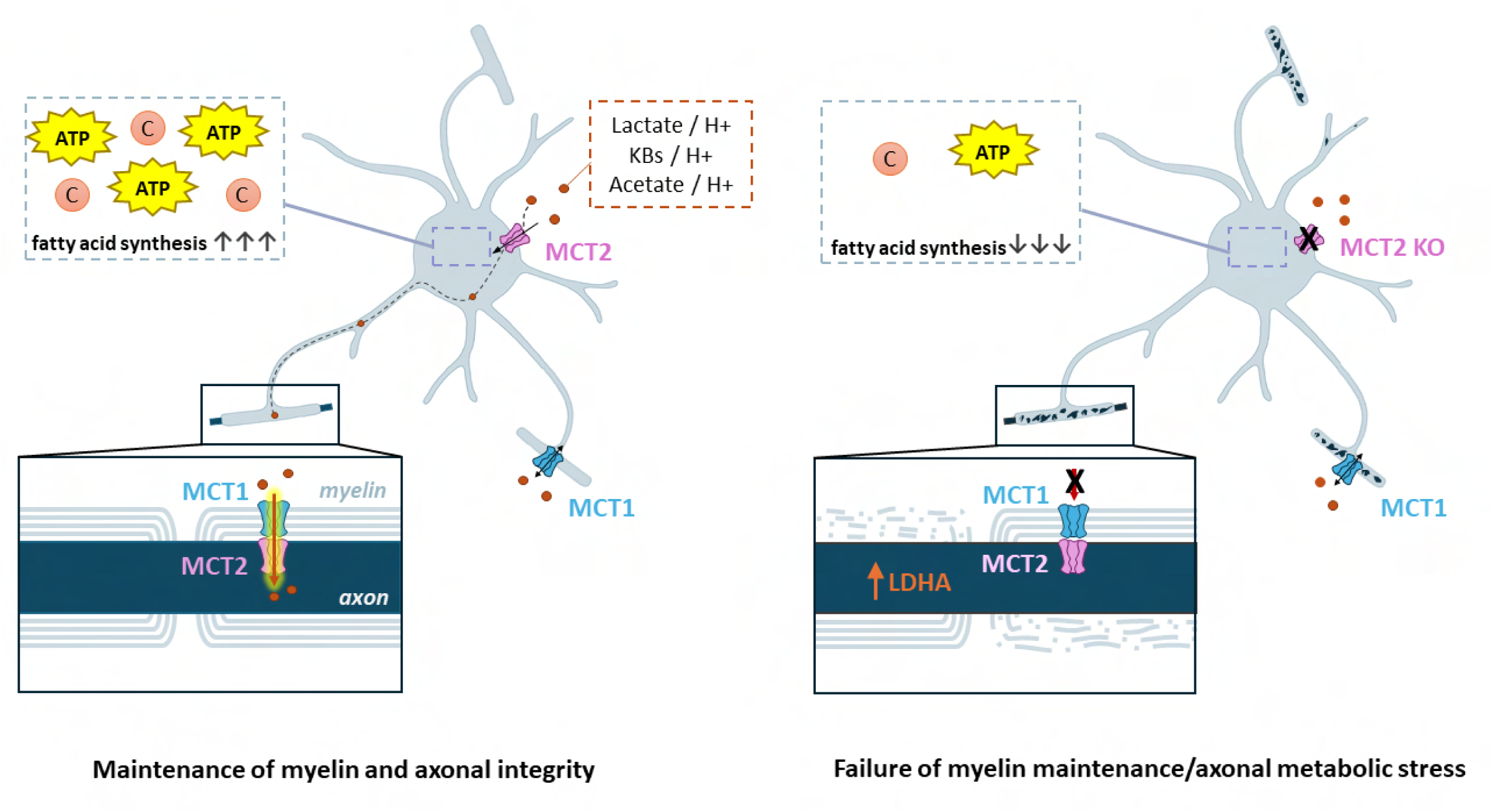

## INTRODUCTION

Metabolic alterations are a well-recognized feature of many, if not all, neurodegenerative disease(*1*). These include multiple sclerosis (MS)(*2–5*), an inflammatory demyelinating disease of the central nervous system (CNS) that represents the leading cause of acquired non-traumatic disability in young adults(*6*). While current treatments effectively treat symptoms associated with inflammatory bouts during relapsing-remitting stages of MS, preventing disease progression that results into permanent neurological handicap remains a major challenge(*7*).

Energy deficit observed in the CNS of patients suffering from neurodegenerative diseases is thought to significantly contribute to brain dysfunction, as the brain is a major consumer of body’s oxygen and glucose (around 20%) while it represents only a 2% of the body’s weight(*8*). While synaptic activity is considered as the most ATP-demanding process in the CNS(*9*), synthesis of myelin sheaths by oligodendrocytes also imposes a high metabolic cost(*10*). Myelinating oligodendrocytes require metabolic fuels to sustain lipid and protein synthesis to ensure myelin maintenance, but also to provide metabolic support to the axons(*11*). Because of high metabolic demands of oligodendroglia, the energy deficit described in MS and other neurodegenerative diseases may significantly compromise the function of these cells, including myelination and the support of axonal integrity, as oligodendrocytes and myelin are highly vulnerable to energy deprivation(*12*). Thus, elucidating metabolic pathways that underlie oligodendroglial function and how these might be affected in MS is crucial to develop therapies to prevent myelin alterations/loss and neurodegeneration, thus disease progression.

Monocarboxylates (MCs), a family of molecules that comprises lactate, pyruvate, acetate, and ketone bodies among others, are increasingly recognized as important metabolic fuels for the CNS(*11, 13–16*). Monocarboxylate trafficking into/out of cells is mediated by monocarboxylate transporters (MCTs). MCTs 1–4 execute the proton-linked transport of MCs across the plasma membrane. MCT1,2, and 4 are expressed in the brain, and out of these, MCT2 has the highest affinity(*17*). This transporter plays an important role in synaptic function, the most energy-consuming process in the brain, as it allows neurons to import astrocyte-derived lactate to be used as energy fuel in the process known as astrocyte-neuron lactate shuttle (ANLS)(*18*). Interestingly, gene expression studies on CNS cells of both mice and humans have shown that the expression of *Slc16a7*, the gene that encodes MCT2, is not restricted to neurons as it is also detected in oligodendroglial cells, including mature oligodendrocytes, and in microglia(*19–23*).

Because of its high metabolic cost, similarly to synaptic function, oligodendroglial myelination might require not only glucose, but also a substantial input of additional metabolic fuels(*11*). Thus, N-acetyl aspartate (NAA) has been described as an important source of both ATP and carbons for myelination(*24, 25*). Regarding MCs, literature suggests these as an important carbon source of myelin lipids: lactate has been shown as a precursor for oligodendrocytic lipid synthesis *in vitro*(*26*), and ketone bodies are preferentially incorporated into myelin lipids over glucose in young rats(*27*). Yet, the mechanisms that control MC import into oligodendrocytes are not clear. Oligodendrocytes are considered to express predominantly MCT1, the intermediate affinity MCT. The expression of this transporter has been detected mainly on myelin both *ex vivo*(*28*) and *in vivo*(*29*). This transporter has been proposed as the primary mediator of the rescue effect of lactate on myelination under low glucose conditions *ex-vivo*(*28*). However, oligodendroglia-specific knockout of MCT1 had no effects on myelination *in vivo* until old age (*30*) suggesting that either MCs are not required for myelination in young and adult mice, or that other MCTs might be involved.

Based on the findings that *Slc16a7* (MCT2-encoding gene) is expressed in oligodendrocytes, we hypothesized that this high affinity monocarboxylate transporter might contribute to the metabolic fitness of these cells, enabling them to maintain myelin sheaths and sustain axonal integrity. Here, we first demonstrated protein expression of MCT2 on myelinating oligodendrocytes in mice and humans. Next, we investigated potential changes in both gene and protein expression of this transporter in MS and showed that it is decreased even in the normally appearing white matter (NAWM) of patients as compared to control subjects. We then performed loss-of-function studies *in vivo* and observed that loss of MCT2 in oligodendrocytes leads to demyelination in absence of oligodendrocyte death, associated with a decreased expression of lipid synthesis-associated enzymes fatty acid synthase (FASN) and acyl-coenzyme A synthetase short-chain family member 2 (ACSS2). Furthermore, axonal damage was observed, coincident with increased axonal expression of lactate dehydrogenase A (LDHA). Finally, pathological consequences of MCT2 deletion were attenuated by provision of ketogenic diet. We conclude that MCT2 enables oligodendrocytes to efficiently import monocarboxylates to support both the high cost of lipid synthesis for myelin maintenance and provision of axons with metabolic fuels. We also suggest that MCT2 loss can be compensated by other, lower-affinity, transporters by increasing extracellular ketone body concentrations via ketogenic diet. Thus, our results provide a novel insight into oligodendrocyte metabolism that could be exploited to design metabolic therapies to enhance oligodendroglial function in diseases such as MS.

## RESULTS

### MCT2 is expressed by oligodendrocytes in mice and humans

As mentioned above, the expression of *Slc16a7* (the MCT2-coding gene) has been reported in human and mouse oligodendrocytes. Thus, we first aimed to investigate whether this mRNA expression is also translated into MCT2 protein expression by performing immunohistochemical analyses on both mouse and human CNS tissue. To investigate the expression by mouse oligodendrocytes we performed co-labelling studies using oligodendrocyte transcription factor 2 (Olig2) as oligodendroglial marker, adenomatous polyposis coli protein (APC)/CC1 as a marker of mature oligodendrocytes, and 2 commercial antibodies for MCT2. Moreover, some labellings were also performed by a previously validated home-made antibody(31). In all cases, MCT2 positive cells were observed in the gray and the white matter, both in the spinal cord (Fig.1) and the brain (Fig.S1). In the gray matter (gm), MCT2+ cells were identified as APC/CC1+ satellite oligodendrocytes (47.78±6.42%), while the rest were NeuN+ neurons (Fig1. A-C,H-I,O). MCT2 positivity was also observed on dendrites and axons (Fig.1H-J). In the white matter (wm), MCT2 expression was detected on axons, as expected, but also on cells that were identified as oligodendrocytes by APC/CC1 expression (Fig.1D-G, K-M). 86.68± 3.10% of the spinal WM oligodendrocytes were positive for MCT2 (Fig.1N), which constituted the 83.28 ± 5.74 % of MCT2 expressing cells in the white matter (Fig.1O). The remaining MCT2+ cells in the wm were identified as Olig2+APC/CC1-cells (Fig.S1A-D), indicating that they were oligodendrocyte progenitors (OPCs). We also investigated MCT2 expression on post-mortem sections of human cerebellum. To label MCT2 we used a previously validated antibody(*32*). As the highest *Slc16a7* transcript expression is detected on neurons(*33*), we first verified whether our antibody labels neuronal cells. Indeed, and similarly to the results obtained in rodent cerebellum(*34*), we observed robust MCT2 positivity in the molecular and granular cell layers, specifically on neurons in the Purkinje cell layer (PCL)(Fig.S2 A-C), and the glomeruli in the granule cell layer (GCL)(Fig.S2D-F). Strong MCT2 positivity was detected on the rosettes, structures that consist of post-synaptic granule cell dendrites and pre-synaptic terminals of mossy fibers (Fig.S2Fi), consistent with previous observations that neuronal MCT2 in mice is enriched at synapses(*34*). After having confirmed the validity of the antibody in labelling neurons, we investigated MCT2 expression in the white matter in both control subjects and patients with MS. In both, we also observed MCT2 positive cells in the cerebellar white matter. Co-labelling with human oligodendroglia marker SOX10, revealed that these MCT2+ cells were oligodendrocytes (Fig.2A-F, Fig.2G-I). In addition, these cells were frequently organized in triplets, typical of myelinating oligodendrocytes (Fig.2A-C). SOX10+ cells in the GCL also expressed MCT2, although those in the WM appeared to be labelled more strongly (Fig. S2G-I). Thus, MCT2 is expressed by myelinating oligodendrocytes throughout the mouse CNS, and in the human cerebellar white matter.

**Figure 1.**
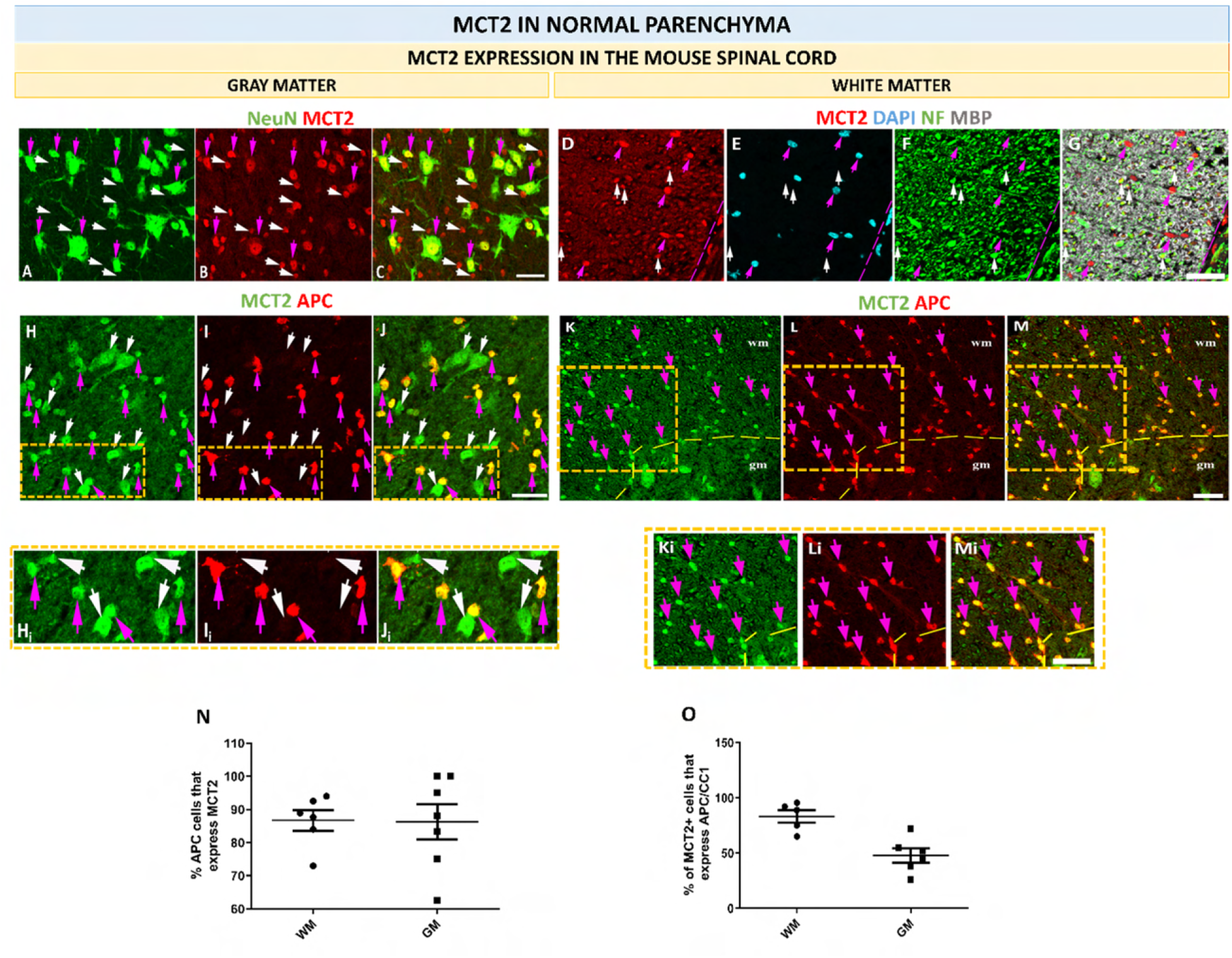
Oligodendrocytes express MCT2 in the mouse spinal cord. **A-C**. Co-labelling for NeuN (green) and MCT2 (red) in the spinal cord gray matter show that MCT2 is seen on neurons (NeuN+ MCT2+, purple arrows) and MCT2+ NeuN-cells (white arrows). **D-G**. Co-labelling for MCT2 (red), DAPI (blue), neurofilament (NF, in green) and MBP (gray) in the mouse spinal cord white matter show that MCT2 is expressed on cells indicated by purple arrows (MCT2+ DAPI+ NF-) and on axons, characterised by MCT2+ NF+ staining and indicated by white arrows. **H-J**. Co-labelling for MCT2 (green) and APC (red) in the gray matter of the spinal cord. MCT2 is seen on neurons (MCT2+ APC-, white arrows) as well as on MCT2+ APC+ oligodendrocytes, including satellite oligodendrocytes, indicated by purple arrows. **Hi-Ji**. Higher magnification of the areas indicated by orange dashed rectangle in H to J. **K-M**. Co-labelling for MCT2 (green) and APC (red) in the mouse spinal cord white matter reveal MCT2+APC+ oligodendrocytes, indicated by purple arrows. **Ki-Mi**. Higher magnification of the areas indicated by dashed orange rectangles in K-M. Scale bars=20µm. NeuN= Neuronal Nuclei, NF= neurofilament, APC=Anti-Adenomatous Polyposis Coli, DAPI=4’,6-diamidino-2-phenylindole; MBP=myelin basic protein, wm= white matter, gm= gray matter. **N.** Quantification of the percentage of APC+ oligodendrocytes that express MCT2 in the spinal cord white and gray matter of 4–5-month-old mice. **O**. Quantification of the percentage of MCT2+ cells that express mature oligodendrocyte marker APC/CC1 in the spinal cord. n=images of separate WM and GM areas from at least 3 different mice. WM= white matter, GM= gray matter.

**Figure 2.**
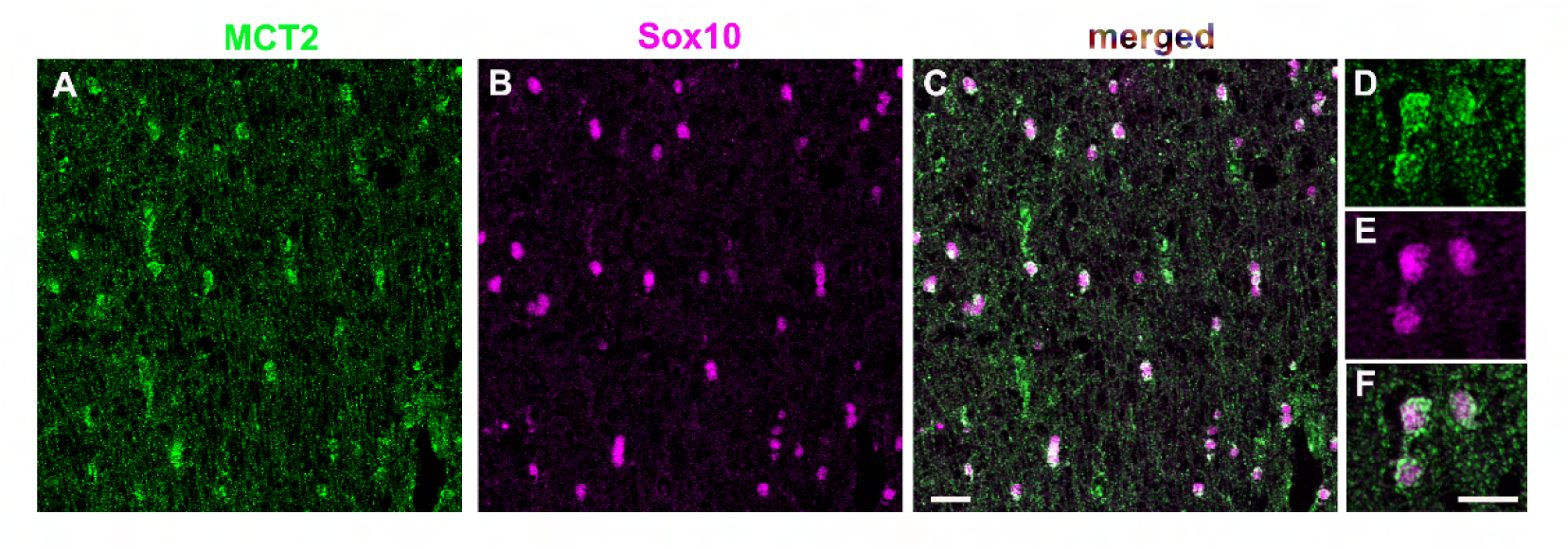
Cerebellar normal appearing white matter (NAWM) from a patient with MS. **A.** Labelling for MCT2 (green). **B.** Labelling for Sox10 (purple), marker of oligodendroglia. **C.** Co-labelling for Sox10 and MCT2. **D-F.** High power view images of MCT2 expressing Sox10+ cells in the NAWM. Similar cellular expression pattern is observed in non-neurological controls. Scale bars=20µm in C, 10 µm in F.

### MCT2 gene and protein expression changes in MS

We then investigated whether MCT2 gene and protein expression by oligodendrocyte were altered in MS.

#### Gene (Slc16a7) expression changes

The analyses of *Slc16a7* expression changes in MS was performed *in silico* using single nucleus RNA-seq data published by Trobisch et al. 2022(*22*). Among the identified cell types, these authors defined four clusters of oligodendrocytes. The two of these were primarily composed of control oligodendrocytes and defined as homeostatic clusters SLC5A11 and LINC01608, while the other two, containing high proportions of reactive oligodendrocytes and defined as clusters HSPA1A and SGCZ, were amplified in MS (Fig.3A). We first detected *Slc16a7* expression across the 4 clusters, after which we compared levels of expression among the clusters. We observed that the two homeostatic clusters present higher expression than the two reactive ones, with significant differences detected between SLC5A11 and the rest of the clusters (Fig. 3B; adjusted p values SLC5A11 vs LINC01608: 0.002, SLC5A11 vs HSPA1A: 0.0162 and SLC5A11 vs SGCZ: 4.06e-8). Interestingly, SLC5A11 was validated in the original publication as the cluster predominantly associated with control white matter. We then investigated potential changes in *Slc16a7* expression in patients with MS versus controls. In the SLC5A11, the homeostatic cluster enriched in the white matter, *Slc16A7* expression significantly decreased in patients with MS (Fig.3C; p value: 0.034), whereas the comparison of general oligodendrocyte *Slc16a7* expression levels (all clusters combined) showed only a tendency for a decrease in the MS group (not shown). As there was a large age gap within both control and MS groups (35-88 years), the subjects were then stratified by age, with the cut-off at 65 years (that allowed us to have sufficient subjects in both “young” and “old” groups, for both controls and patients with MS). In subjects younger than 65 years, *Slc16a7* mRNA levels were similar between controls and MS patients (adjusted p value: 0.509). However, with age, MS patients older than 65 years showed a significantly decreased *Slc16a7* expression levels compared to controls older than 65 (Fig.3D; adjusted p value: 0.0406). Moreover, because our analyses included oligodendrocytes from different regions (brain vs spinal cord), and regional differences in gene expression among different human oligodendrocyte populations have been documented(*35*), we then re-analysed *Slc16a7* expression by oligodendrocytes from the cortex and the spinal cord, including age as a control variable in the statistical models, in order to investigate whether these regions were differentially affected. Interestingly, while in the cortex we observed only a non-significant tendency for a decrease, in the spinal cord, in which oligodendrocytes maintain longer internodes(*36*) thus have larger synthetic needs, a larger decrease in *Slc16a7* expression was detected (Fig.3E-F; p value cortex: 0.66, p value spinal cord: 0.08).

**Figure 3.**
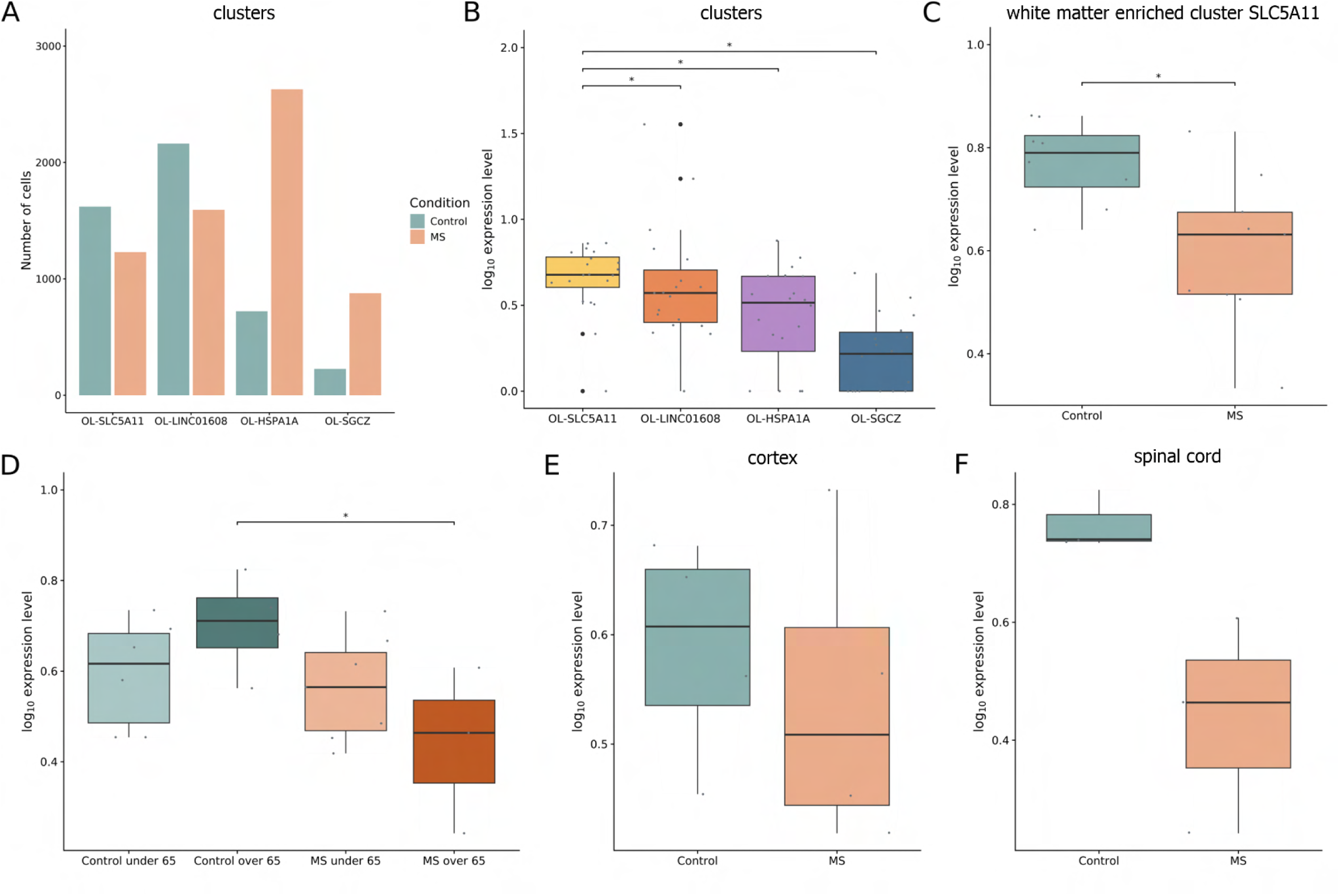
Analyses of Slc16A7 expression changes in progressive MS. **A.** Number of oligodendrocytes (Y-axis) analysed by cluster (X-axis) and condition (bar colour). **B-F.** Normalized and logarithmically scaled Slc16A7 gene expression values across different comparisons: **B**. oligodendrocyte clusters, **C.** SLC5A11 white matter-enriched oligodendrocyte cluster, **D.** stratification by age groups, **E.** cerebral cortex, and **F**. spinal cord. Boxplots represent the median (horizontal line) and interquartile range (box). Whiskers extend to the smallest and largest values within 1.5 times the interquartile. range. Outliers are shown as large black dots, while small gray dots represent individual samples. Statistical significance is indicated by * for p-value (single comparison) or adjusted p-value (multiple comparisons) < 0.05. MS: multiple sclerosis; OL: oligodendrocytes.

Thus, these data suggest a decrease in *Slc16a7* expression by oligodendrocytes in patients with progressive MS that affects preferentially the spinal cord and appears attributable to both the appearance of reactive oligodendrocyte clusters, as well as the decrease in the expression within the homeostatic cluster SLC5A11.

MCT1 is another MCT that imports monocarboxylates, although with a significantly lower affinity than MCT2(*13*). Thus, we also investigated variations in *Slc16a1*, the gene encoding MCT1, using the same analyses as those performed for *Slc16a7.* First, we observed that the expression levels of *Slc16a1* in oligodendrocytes are lower than those of *Slc16a7*. In addition, unlike *Slc16a7* that was enriched in the white matter cluster SLC5A11, *Slc16a1* expression was significantly lower in this cluster as compared to the homeostatic cluster LINC01608 and reactive clusters HSPA1A and SGCZ (adjusted p values= 0.00029, 0.016, and 4.06e-8, respectively) (Fig. S3A). No significant differences in the *Slc16a1* expression within the white matter cluster were detected between controls and MS (Fig.S3B). The analyses with the age cutoff at 65 years showed a non-significant decreasing tendency (p=0.0954) between controls vs patients older than 65 years (Fig.S3C). In addition, while a non-significant increase in *Slc16A7* expression had been observed in the controls with age (Fig.3D), changes in control *Slc16a1* expression with age were not observed (Fig.S3C). Although age-adjusted regional analyses of *Slc16a1* expression changes in oligodendrocytes in MS showed decreasing tendencies, these differences were far from reaching statistical significance both in the cortex (p=0.42; Fig.S3D) and the spinal cord (p=0.28; Fig.S3E). Thus, the pattern of *Slc16A1* expression among human oligodendrocyte clusters is different from that of *Slc16a7*, and the extent of change in progressive MS is much lower.

#### MCT2 protein expression changes in MS

We then examined MCT2 protein expression in post-mortem sections of the human cerebellum in control subjects versus patients with MS. Clinical features of the subjects analysed are presented in Table S1. Images of sections cut from the blocks were first stained with Luxol Fast Blue (myelin staining), cresyl violet (neuronal cell bodies), and major histocompatibility complex II (MHC II; marker for inflammatory cells (not shown) to characterize the tissue. We used Sox10 antibody as a marker for oligodendroglia. SOX10+ and MCT2+ cells were quantified at the rims and lesion cores of chronic and chronic active lesions, and within the normal appearing white matter (NAWM) of MS and non-neurological controls.

Although numbers of SOX10+ cells per area were generally increased in the NAWM of MS cases versus controls (669.2±55.55 cells/mm^2^ in controls and 1325 ±406.2 cells/mm^2^ in MS), the variability among MS cases was important, thus the differences between the two experimental groups were not significant (Fig. 4A). We then investigated the numbers of SOX10+ cells in the lesion rim or perilesion and the lesion core of different MS cases and compared them to those in the corresponding NAWM. The number of SOX10+ cells per area in the perilesion was 1436±575.0, and in the lesion core decreased to 43.52±41.43. Statistical differences were found between the perilesion (rim) and the lesion core (Fig.4B).

**Figure 4.**
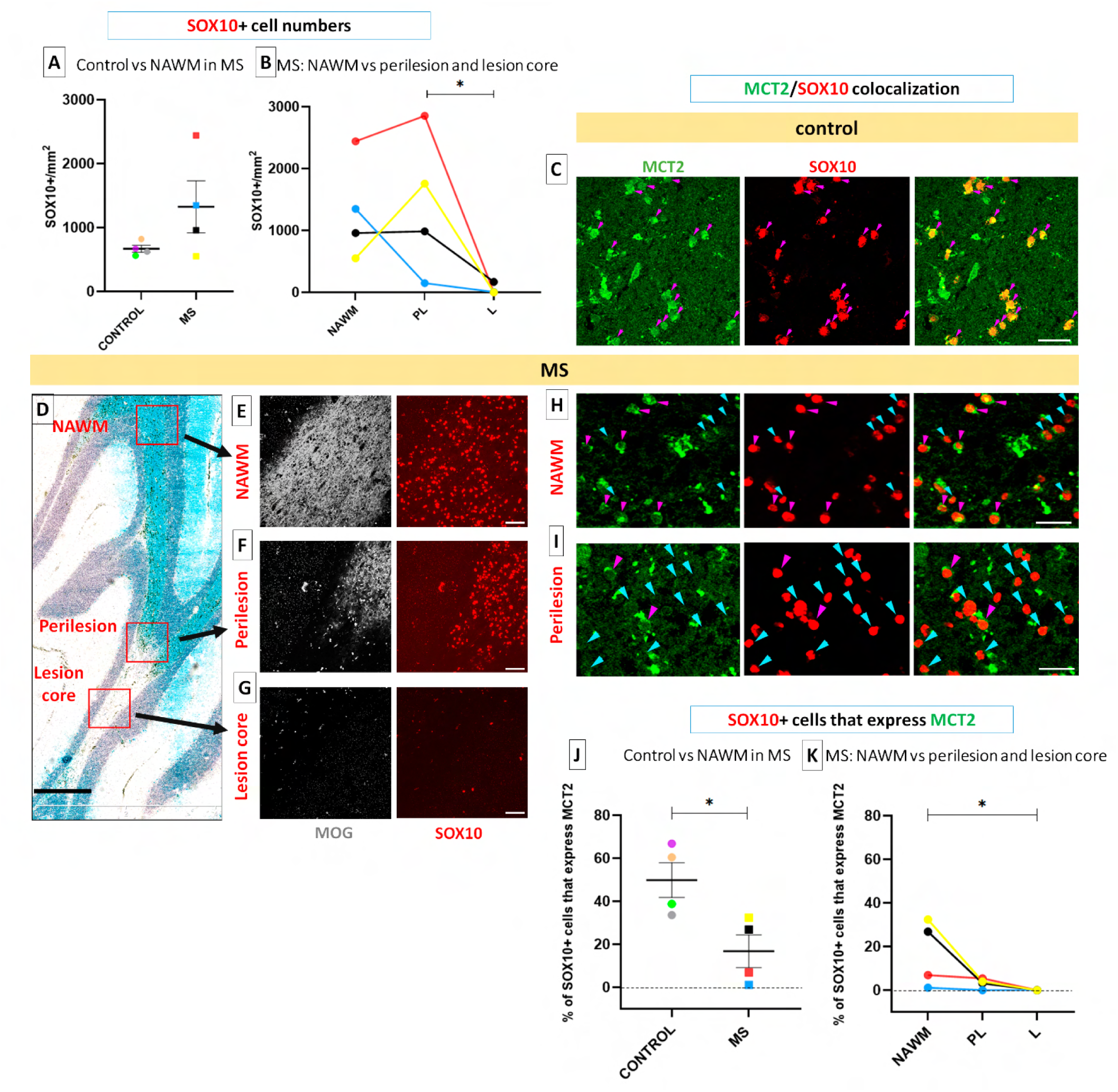
Analyses of SOX10/MCT2 numbers and colocalization in the cerebellum of controls vs patients with MS. **A**. SOX10+cell numbers in MS NAWM vs controls. **B**. Lines represent different patients; in 2/5 patients, SOX10+numbers increase perilesionally, 1/5 shows no change, and in 2/5 SOX10+ numbers decrease. A significant downregulation of SOX10 was observed in the lesion core compared to the rim (perilesion), *p=0.04, One-way ANOVA with Dunn’s multiple comparison. All patients show SOX10+ cell depletion in the lesion core. **C.** Co-immunohistochemistry for SOX10 (red) and MCT2 (green) in the white matter of a control subject. **D**. Luxol fast blue image of cerebellar tissue of the patient with MS harboring a chronic active lesion illustrates different tissue areas analyzed. **E-G**. Co-immunohistochemistry for SOX10 (red) and MOG (gray) in the NAWM (E), perilesion (F), and lesion core (G) areas indicated by squares in D. **H,I** Co-immunohistochemistry for SOX10 (red) and MCT2 (green) in the NAWM (H) and perilesion (I). **J-K**. Quantification of MCT2/SOX10 colocalization. **J**. The % of SOX10+ cells that express MCT2 is the highest in the control subject white matter, decreases significantly in the MS NAWM, Welch’s unpaired t-test (unequal variances) * p=0.0246 (X), and **K**. further decreases towards lesion core; One-way ANOVA with Dunn’s multiple comparisons test indicates a significant decrease between NAWM and the lesion core, *p=0.0240. Controls N=4 and MS patients N=4. **C,H,I** Purple arrows: MCT2+cells; light blue arrows =MCT2-cells. Scale bars: C,H,I= 20 µm, D=2 mm, E,F,G=50 µm. NAWM=normal appearing white matter. PL=perilesion. L=lesion core.

Then, we analysed the co-localization between MCT2 and SOX10, in controls versus in MS NAWM and changes in MS tissue according to the lesion proximity (Fig.4C-K). Interestingly, the percentage of SOX10+ cells that express MCT2 decreased from 49.92±8.093% in controls to 16.84±7.566% in NAWM MS (p=0.0246) (Fig. 4J). Moreover, this percentage in patients with MS further decreased to 3.133±1.145% in the rim (perilesion) and to 0% in the lesion core. A statistically significant difference was detected between the NAWM and lesion core.

Thus, MS patients show a decrease in MCT2 expression by oligodendroglial cells in the NAWM compared to the controls, and this expression becomes negligible in the lesion proximity (perilesion), although oligodendroglial cells are present in significant numbers. Importantly, chronic lesion core contains very low numbers of SOX10+cells that do not express MCT2.

Altogether, we conclude that MCT2 is expressed by myelinating oligodendrocytes throughout the mouse central nervous system and in the white matter of the human cerebellum, and that this expression decreases in progressive MS.

### MCT2 deletion in myelinating oligodendrocytes in the spinal cord leads to demyelination

MCT2 is a high affinity transporter for lactate and ketone bodies. These molecules have been associated with lipid synthesis in oligodendrocytes(*26, 27, 37*). Thus, we hypothesized that MCT2 may be important for lipid synthesis and myelin maintenance. To investigate the role of MCT2 in myelin maintenance, we aimed to delete the *Slc16a7*, the gene coding for MCT2, in myelinating oligodendrocytes. For that, we used MCT2^lox/lox^ mice(*38*) and the oligodendrotropic AAV, Olig001, with capsid tropism for myelinating oligodendrocytes(*39–41*). As spinal cord oligodendrocytes maintain long internodes(*36*), and we detected a preferential *Slc16a7* decrease in the spinal cord in MS (Fig.3F), we tested the effect of S*lc16a7* deletion in the spinal cord oligodendrocytes. Thus, we injected the Olig001 construct carrying the Cre recombinase tagged with GFP (Olig001-Cre-GFP) or the control Olig001-GFP AAV into wildtype (wt) or MCT2^lox/lox^ mouse dorsal funiculus (Fig.5A). Mice were sacrificed at 3 weeks post injection, to allow adequate AAV expression and recombination. GFP expression in the white matter was observed in both groups. Transduced cells were Olig2+ (62.19±14.86%)(Fig.S4D-F) and APC/CC1

**Figure 5.**
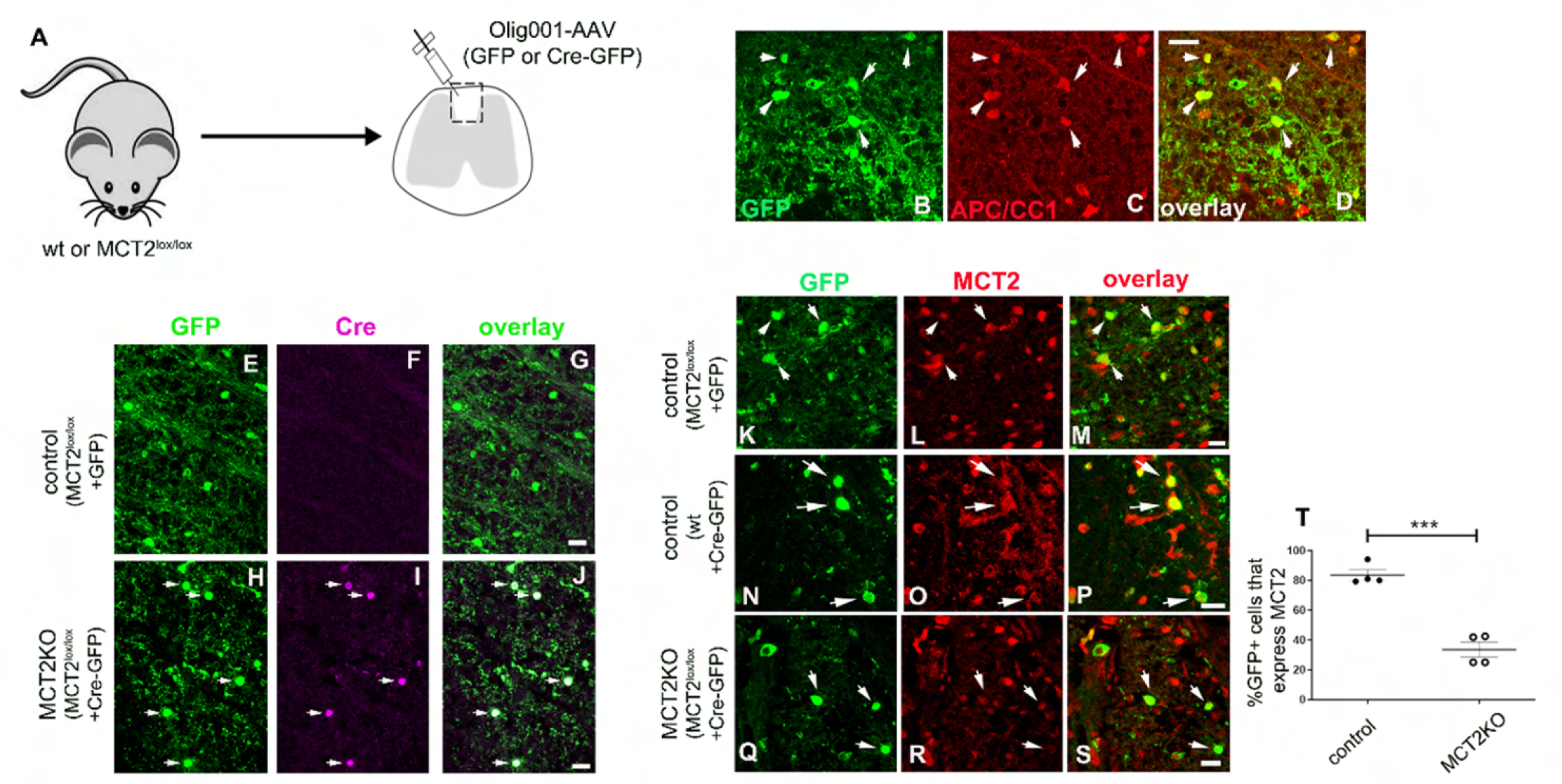
MCT2 deletion in myelinating oligodendrocytes using AAV-mediated Cre-Lox approach. **A.** Schematic presentation of the experimental strategy employed to delete Slc16a7 in mature oligodendrocytes. Wildtype (wt) and MCT2^lox/lox^ were injected with oligodendrotropic Olig001-AAV carrying either GFP or Cre-GFP construct in the spinal cord dorsal white matter. Mice were sacrificed at 3- and 6-weeks post injection. **B-D**. Co-immunolabelling for GFP in green (B) and mature oligodendrocyte marker APC/CC1 in red (C). **D**. Overlay of B-C. Arrowheads indicate double-labelled cells. **E-J**. Co-labelling for GFP in green (E,H) and Cre-recombinase in magenta (F,I) in MCT2^lox/lox^ mice injected with Olig001-GFP (E-G) or Olig001-Cre-GFP (H-J). **G**. Overlay of E-F. **J**. Overlay of H-I. Arrowheads indicate double-labelled cells, only apparent in Olig001-Cre-GFP injected MCT2^lox/lox^ mice. **K-S**. Co immunolabelling for GFP in green (K,N,Q) and MCT2 in red (L,O,R) **M**. Overlay of K-L. **P**. Overlay of N-O. **S**. Overlay of Q-R. White arrows indicate GFP+ cells. **K-M**. MCT2^lox/lox^ mice injected with Olig001-GFP. **N-P**. Wt mice injected with Olig001-Cre-GFP. **Q-S**. MCT2^lox/lox^ mice injected with Olig001-Cre-GFP (MCT2KO mice). **T**. The proportion of GFP+ cells that are MCT2+ significantly decreases in MCT2KO compared to controls. Unpaired student t-test, (*) p=0.0007. n=4 mice per group. Scale bars=10µm.

+(76.42±16.38%) (Fig.5B-D), demonstrating their identity as oligodendroglia and post-mitotic oligodendrocytes, respectively. GFP label never colocalized with IBA1, a microglia/macrophage marker (Fig. S4G-I). Transduced cells were also negative for astrocyte marker GFAP (Fig.S4J-L), except for an occasional cell (less than 1%) in some animals. We also verified the expression of Cre-recombinase using Cre-specific antibodies and observed co-expression with GFP in Olig001-Cre-GFP injected, but not Olig001-GFP injected animals (Fig.5E-J). Thus, using this experimental paradigm, we successfully expressed Cre in mature oligodendrocytes in the white matter that we could follow by monitoring GFP expression.

We also observed variable degrees of transduction in gray matter (Fig.S4B-C), likely due to AAV spreading. In the gray matter, transduced cells were identified as APC/CC1+ oligodendrocytes, but also as APC/CC1-cells, of neuronal morphology (not shown).

We then investigated whether MCT2 protein expression was diminished in recombined cells. We performed co-immunolabellings for MCT2 and GFP. MCT2 expression was observed on GFP+ cells, indistinguishably from GFP-cells in the white matter of MCT2^lox/lox^ mice injected with Olig001-GFP AAV (Fig.5K-M), as well as in wt mice injected with Olig001-Cre-GFP (Fig.5N-P). Yet, very weak or absent MCT2 labelling was observed on many GFP+ cells in MCT2^lox/lox^ mice injected with Olig001-Cre-GFP, while MCT2 expression was present on GFP-cells (Fig.5Q-S). The proportion of cells positive for MCT2 among the GFP+ population decreased from 83.60± 3.52 in MCT2^lox/lox^ mice injected with Olig001-GFP (from herein referred to as controls) to 45.07± 12.09 in MCT2^lox/lox^ mice injected with Olig001-Cre-GFP (from herein referred to as MCT2KO) (Fig.5T). Thus, Olig001-Cre-GFP transduction in MCT2^lox/lox^ mice successfully decreased levels of MCT2 in transduced cells.

We then investigated the evolution of axonal and myelin protein expression following MCT2 deletion in white matter oligodendrocytes. For this, we generated an additional cohort of control vs MCT2KO mice that we sacrificed at 6 weeks after injection. Co-immunolabellings for GFP, SMI31 (phosphorylated neurofilament) and myelin basic protein (MBP) revealed transduced oligodendrocytes surrounded by myelinated axons in the *controls* at both 3 and 6 weeks (Fig. 6A-D). Yet, in the MCT2KO, GFP+ cells were frequently observed within the areas of myelin loss, already evident at 3 weeks, and still present at 6 weeks (Fig.6E-H). Quantification of MBP+ area fraction revealed a significant decrease in MCT2KO injected white matter at both time points, decreasing those values from 63.98± 7.58% to 38.85±2.98% at 3 weeks and from 61.84±1.78% to 32.90±1.89% at 6 weeks (Fig.6I).

**Figure 6.**
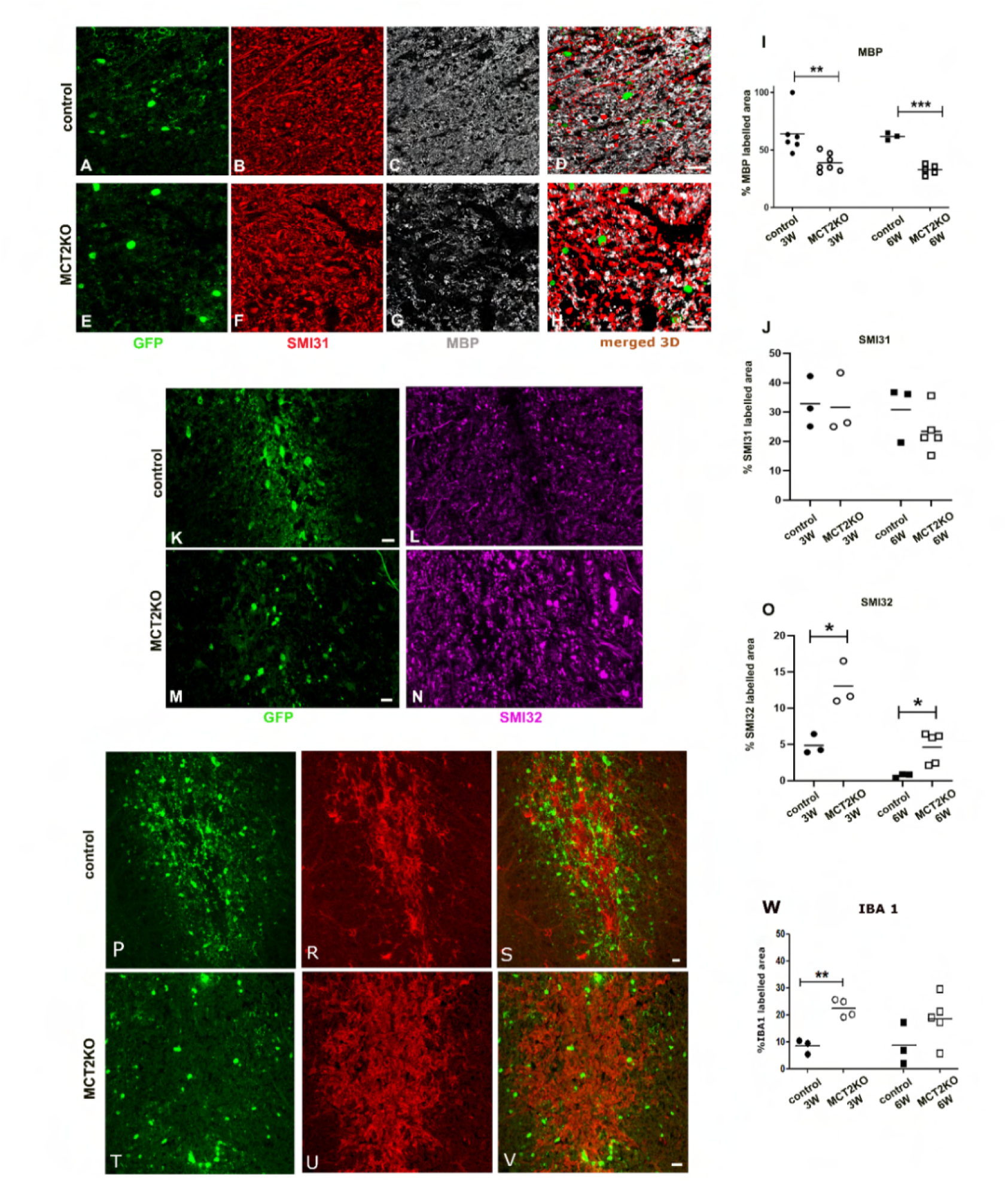
Demyelination, axonal injury and microgliosis in MCT2 KO mice. **A-G.** Co-labelling for GFP in green (A,E), SMI31 in red (B,F), and MBP in grey (C,G) in control (A-C) and MCT2KO mice (E-G) at 6 weeks post Olig001-AAV injection. **D**. 3D overlay of A-C. **H**. 3D overlay of E-G. **I**. The percentage of MBP-labelled area within the total WM area significantly decreases in MCT2KO at 3 weeks; Student unpaired t-test (**) p= 0.0074, n=6 controls vs 7 MCT2 KO; as well as at 6 weeks; Student unpaired t-test (***) p<0.0001, n= 3 controls vs 5 MCT2 KO. **J**. No significant changes in the percentage of SMI31-labelled area between the groups. n=3 for all cases except for 6 weeks Cre-GFP, where n=5. **K-N.** Co-labelling for GFP in green (K,M) and SMI32 in magenta (L,N) in control (K-L) and MCT2KO mice (L-N) at 6 weeks post Olig001-AAV injection. **J.** White matter area fraction labelled with SMI32 is significantly increased in MCT2KO at 3 weeks [(*) p= 0.0272 (Welch’s t test)] and 6 weeks after Olig001-AAV injection [(*) p= 0.0357 (Mann-Whitney test)]. **P-V.** Co-labelling for GFP in green (P,T) and IBA1 in red (R,U) in control (P-R) and MCT2KO mice (T-U) at 3 weeks post Olig001-AAV injection. **S**. Overlay of P-R. **V**. Overlay of T-U. **W**. The percentage of Iba1 labelled area within the total white matter area is significantly increased at 3 weeks post Olig001-AAV injection in MCT2KO mice, Student unpaired t-test, (**) p=0.0018, but not at 6 weeks, p=0.1244. n=3 for controls at both time points, n=4 for MCT2KO at 3 weeks and n=5 for MCT2KO at 6 weeks. Scale bars=10µm.

### Axonal injury in MCT2KO mice

We then investigated axonal integrity by performing immunolabellings for SMI31 (phosphorylated neurofilament) and SMI32 (non-phosphorylated neurofilament). Although we observed slight disorganization of SMI31 staining in MCT2KO mice (Fig.6B,F), quantification of percentage area labelled with SMI31 revealed no significant differences between the groups neither at 3 nor at 6 weeks (Fig. 6J). We then assessed axonal damage by performing immunohistochemistry for SMI32, associated with axonal pathological states(*42*). The percentage of the area labelled with anti-SMI32 significantly increased in MCT2KO mice both at 3 and 6 weeks (controls 4.87±0.79%, MCT2KO 13.05±1.74%, *p = 0.027 at 3 weeks; controls 0.70±0.16% vs MCT2KO 4.62±0.96% , *p = 0.0357, 6 weeks), and swollen axons strongly positive for SMI32 were evident in MCT2KO but not the control mice (Fig. 6K-O).

Next, we investigated inflammation in MCT2KO mice. We performed labellings for Iba1+ cells to investigate presence of myeloid cells, in control vs MCT2KO mice both at 3 and 6 weeks post Olig001-AAV injection. Iba1+ (myeloid)(Fig.6R,U) and CD45+ (hematopoietic/activated microglia) cells (not shown) were present in the white matter areas of both control and MCT2KO mice. The percentage of white matter area occupied by Iba1+ staining significantly increased in MCT2 KO mice at 3 weeks (8.423 ± 1.538% in controls, 22.46 ± 1.651% in MCT2KO, **, p = 0.0018, Fig.6R,U,W). The increase persisted but was no longer significant at 6 weeks (8.65±4.53% in controls and 18.87±3.50% in MCT2KOs, Fig.6W). In addition, quantification of CD3+ cells (T cells) at 6 weeks revealed a trend towards an increase in the MCT2KO (56.37±45.01 CD3+ cells/mm^2^ for control and 138.80±52.21 CD3+ cells/mm^2^ for MCT2 KO). Thus, Olig001-AAV transduction itself in the white matter leads to a low degree of inflammation, which is upregulated in MCT2KO, particularly at early time points.

To gain a better insight into white matter changes in MCT2KO, we performed ultrastructural analyses using transmission electron microscopy (TEM). To ensure that the analyses were performed in the transduced white matter area, we combined TEM with immunogold labellings for GFP in control vs MCT2KO mice sacrificed at 6 weeks post AAV injection. These analyses revealed that in controls, oligodendrocytes, identified by their immunogold labelling for GFP and Olig2, exhibited a typical rounded morphology, dark spherical nucleus, with some small chromatin clumps, slightly dense cytoplasm, with abundant ribosomes and short cisternae of rough endoplasmic reticulum (ER) (Fig.7A) and were surrounded by thick myelin-enveloped axons (Fig.7B). Except for the small areas surrounding the needle tract, in which evidence of complete remyelination (thin myelin) was evident, the myelin status in these animals appeared largely normal, and myelin-debris-filled macrophages were absent. Conversely, in MCT2KOs, oligodendrocytes displayed an altered/reactive morphology, with lighter nuclei and numerous mitochondria (Fig.7C), but persistence of short cisternae of the ER (Fig.7Ci). Of note, it was confirmed that these cells were indeed oligodendrocytes as they were positive for Olig2 (Fig.S5). Demyelinated axons were visible in the vicinity of these cells (Fig.7C-D), as well as microglial cells, some of which had many lipid droplets within their cytoplasm indicating they had ingested myelin debris (Fig.7D-arrows). Microglial cells were also observed in contact with myelin sheaths that appeared separated from the axon as an empty space (Fig.7F), indicating myelin disintegration. Besides demyelinated axons, we also observed axons with very thin myelin sheaths (Fig.7G), demyelinated axons showing accumulation of electron-dense bodies indicative of axonal damage (Fig.7G arrow), and occasional evidence of axonal degeneration reflected by the presence of myelin sheaths with axonal debris inside (Fig.7H).

**Figure 7.**
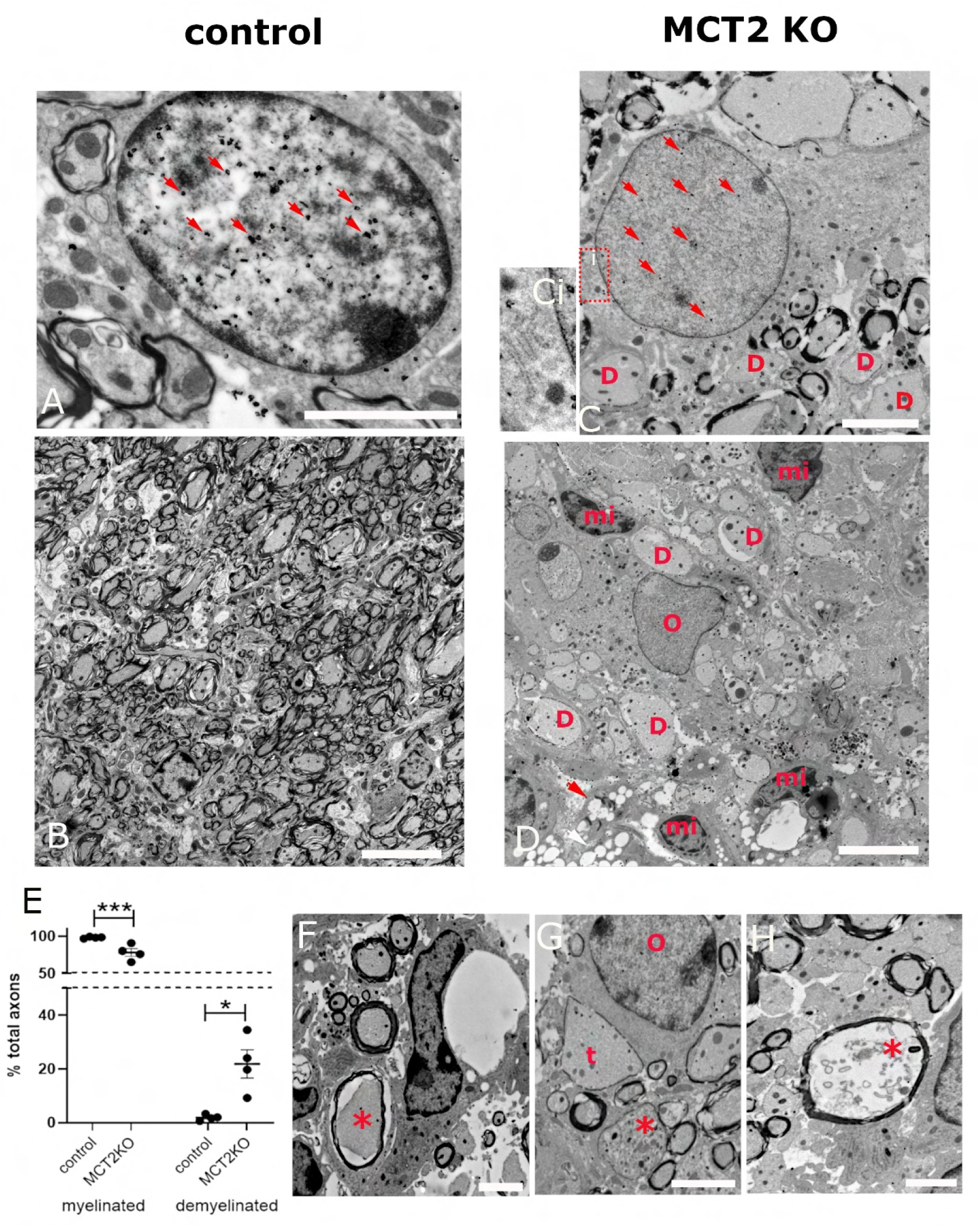
Immunogold labelling for GFP combined with transmission electron microscopy shows ultrastructural alterations of oligodendroglia, myelin, and axonal integrity in MCT2KO mice. **A.** GFP+ cell in the control mouse surrounded by myelinated axons. **B.** Low power images reveal oligodendrocytes and myelinated axons in the control mouse. **C.** GFP+ cell in a MCT2KO mouse next to demyelinated axons (D). Inset **Ci** shows short cisternae of rough ER, typical of an oligodendrocyte. **D**. Low power images of a MCT2KO mouse show demyelinated axons (D), numerous microglial cells (mi) and myelin debris filled microglia/macrophages (arrows). **E.** Quantification of myelinated vs demyelinated axons in controls and MCT2KOs. The percentage of myelinated axons is significantly decreased in MCT2KOs (*) p=0.03, while that of demyelinated axons is significantly upregulated in MCT2KO (*) p=0.0308. Unpaired Welch’s t-test. For controls n=4 and for MCT2KO n=4. **F-G**. TEM images of MCT2KO mouse dorsal funiculus. **F**. An axon (*) on which myelin sheath is separated by an empty space, potentially indicating detachment, in close contact with microglial cell. **G**. A demyelinated axon with accumulation of electron dense bodies, indicative of axonal injury (*). An axon with thin myelin is also observed (t). **H**. Axonal degeneration reflected by a myelin sheath that contains debris, but not an axon. Scale bars A-C, F-G=2µm, B,D=5µm.

We then quantified myelinated and demyelinated axons throughout the dorsal funiculus in which we detected GFP labelling in control vs MCT2KO mice. The percentage of myelinated axons significantly decreased from 98.11±0.54% in the controls to 78.13±5.23% in the MCT2KOs. Conversely, the percentage of demyelinated axons significantly increased from 1.89±0.54% in the controls to 21.87±5.23% in the MCT2KOs (Fig.7E).

Thus, MCT2 deletion in white matter oligodendrocytes leads to demyelination, axonal damage, and altered oligodendroglial morphology.

### Deletion of MCT2 in spinal cord neurons does not lead to demyelination

Although the transduction in the white matter was restricted to mature oligodendrocytes, in the neighboring gray matter we also observed transduced neurons. While these neurons do not project their axons in the dorsal funiculus (thus excluding axonal effects on myelin integrity in that area), it could be that MCT2 deletion in these neurons could alter extracellular environment and induce demyelination in the neighboring white matter. To investigate whether MCT2 deletion in spinal cord neurons leads to demyelination of the neighboring white matter, we deleted MCT2 specifically in these neurons and not oligodendrocytes. We injected the AAV carrying Cre-GFP or the hSyn-GFP (control) under neuron-specific hSyn promoter, that was previously used to study the effect of MCT2 deletion on hippocampal neurons(*38*), in the spinal cord of MCT2^lox/lox^ mice using the same injection settings as those used for Olig001-AAV injection (Fig.S6A). As expected, GFP expression was observed only on NeuN+ cells in the gray matter, and no GFP expression was observed in the white matter (Fig.S6B-C), confirming previously reported neuronal specificity. We then analyzed the expression of SMI31 (axonal marker)(Fig.S6E,H), MBP (myelin marker)(Fig.S6F,I) and Iba1 (microglia/macrophage marker)(Fig.S6J,K) expression in the neighboring white matter. Quantification of MBP+ labelled area (Fig.S6L), SMI31+ staining (Fig.S6M), and Iba1+ labelled area (Fig.S6N) did not reveal differences between the hSyn-GFP- and hSyn-Cre-GFP injected mice, indicating that myelination as well as axonal and microglia numbers status were similar between the two groups.

Thus, demyelination seen in MCT2^lox/lox^ mice injected with oligodendrotropic Olig001-Cre-GFP-AAV is due to MCT2 loss in oligodendrocytes and not neurons.

### Demyelination in *MCT2KO* mice is not due to oligodendrocyte death

In principle, demyelination can occur either because myelinating cells die or because they fail to maintain myelin. We first investigated whether MCT2 deletion leads to oligodendrocyte apoptosis. We performed co-immunolabellings for cleaved caspase 3 (CASP3), a marker of apoptotic cells, GFP, and Olig2, an oligodendroglial marker. We observed CASP3+ cells both in control and MCT2KO mice, mostly at 3 weeks after injection (Fig.S7A-F). Their numbers did not differ between the groups, and markedly decreased at 6 weeks (at 3 weeks, 257±118.8 CASP3+ cells controls vs 216.4±119.2 CASP3+ cells in the MCT2KO; 6 weeks 57.22±23.26 controls group vs 35.52±20.85 MCT2KO; Fig.S7G). Yet, no colocalization was seen between CASP3 and GFP (Fig.S7C,F) suggesting that transduced cells were not the ones undergoing apoptosis. In addition, GFP+ cell numbers were similar between *control* and *MCT2KO* mice both at 3 and 6 weeks (200.4±25.74 in controls versus 206.0±29.64 in MCT2KOs at 3 weeks and 191.20±19.54 in controls versus 192.0±23.51 in MCT2KOs at 6 weeks; Fig.S7H), showing that transduced cells persist in the MCT2KO thus excluding the possibility that MCT2-deficient oligodendrocytes may be undergoing non-apoptotic cell death. We then checked whether non-transduced oligodendrocytes were CASP3+, but did not observe CASP3 expression on Olig2+ cells. Moreover, the total numbers of mature oligodendrocytes (APC/CC1+) were not statistically different between the groups (APC+ cells/mm^2^ for controls 440.0±58.79, and for MCT2KO 314.7±63.37). Thorough characterization of CASP3+ cells identity revealed these were CD45+ hematopoietic cells in both groups (not shown), which is consistent with previous reports showing that infiltrating inflammatory cells undergo apoptosis in the CNS(*43, 44*).

Thus, MCT2 deletion in oligodendrocytes does not lead to oligodendroglial death which strongly suggests that *MCT2KO* oligodendrocytes survive but fail to properly maintain myelin.

### *MCT2KO* oligodendrocytes show reduced expression of enzymes involved in fatty acid synthesis

As our results showed demyelination in absence of oligodendroglial death in *MCT2KO* mice, we hypothesized that *MCT2KO* oligodendrocytes survive but fail to maintain myelin. As myelin is mostly lipids, we focused on the expression of enzymes involved in fatty acid (FA) synthesis. We investigated the expression of fatty acid synthase (FASN), an enzymatic complex dimer that catalyzes the first step of FA synthesis, namely the conversion of acetyl-co-A to palmitate (Fig.8A). We observed extensive FASN expression in the white matter. In *control* mice, the absolute majority of GFP+ cells were also FASN+ (Fig.8B-D,H). However, in the *MCT2KO* mice, numerous GFP+FASN-cells were observed (Fig.8E-G). Quantification of GFP+FASN+ cells showed a significant reduction in the proportion of FASN-expressing cells within the GFP population (89.98±2.32% in control vs 47.57±6.08% in MCT2KO mice at 3 weeks and 86.20±2.03% in control vs 55.65±4.95% in MCT2KO at 6 weeks, Fig.8H).

**Figure 8.**
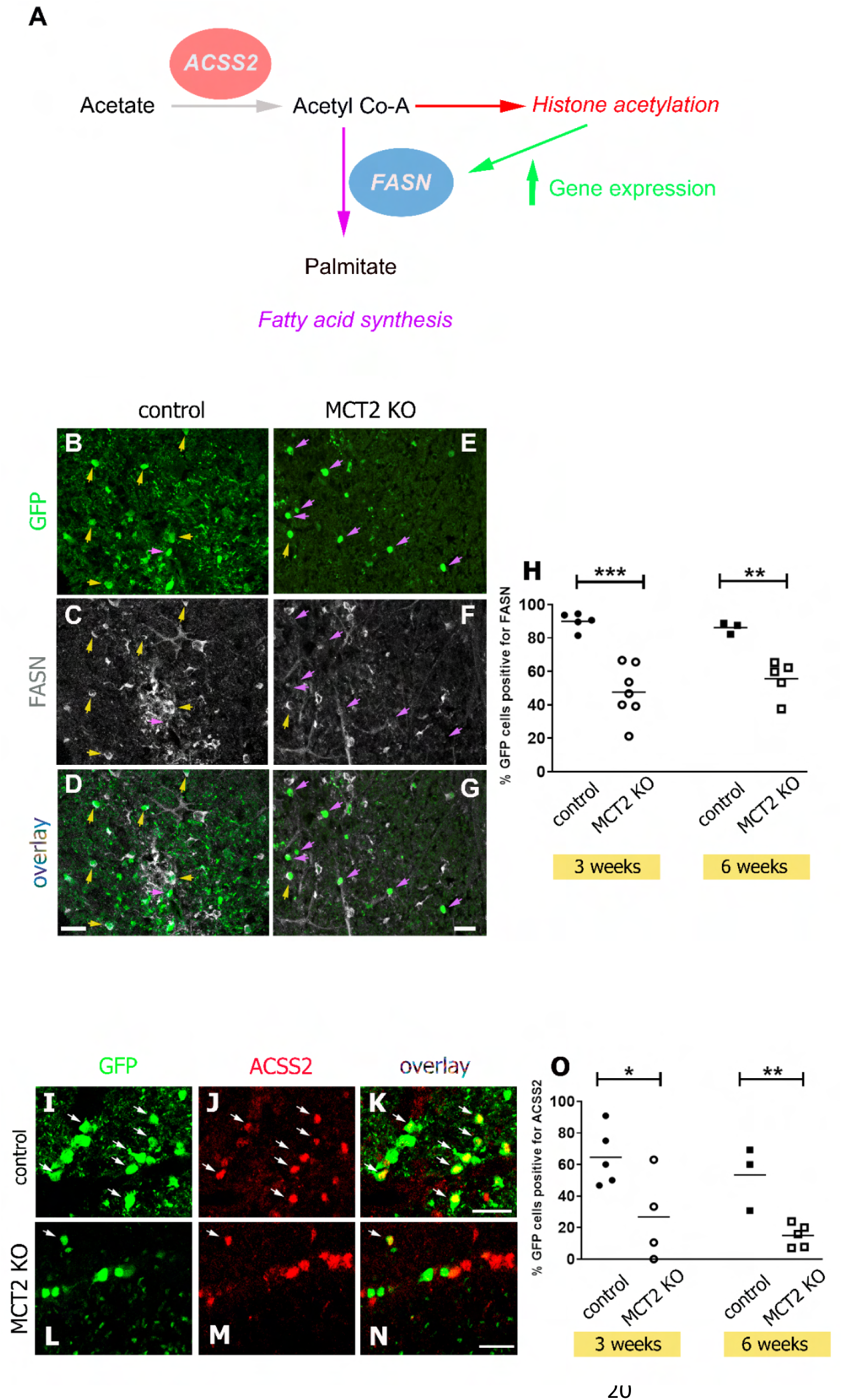
Reduced expression of 2 major enzymes that regulate fatty acid synthesis in MCT2KO oligodendrocytes. **A.** Schematic representation of fatty acid synthase (FASN) and acyl-coenzyme A synthetase short-chain family member 2 (ACSS2) involvement in fatty acid synthesis. ACSS2 regulates lipid synthesis via two distinct, not mutually exclusive mechanism. First, ACSS2 catalyses the conversion of acetate (that can be imported via MCT2) to acetyl co-A. Acetyl Co-A then can be used as a substrate for, on one hand, palmitate synthesis by FASN (1^st^ step of fatty acid synthesis), or on the other hand, for histone acetylation which enhances the expression of the genes that regulate lipid synthesis, including the gene encoding FASN. **B-G**. Co-immunolabelling for GFP in green (B,E) and FASN in grey (C,F). D overlay of B-C. G overlay of E-F. B-D control. E-G MCT2 KO. Yellow arrows indicate GFP+FASN+ cells, purple arrows indicate GFP+FASN-cells. **H**. Quantification of the percentage of GFP+ cell expressing FASN at 3 weeks and 6 weeks. At 3 weeks (***) p = 0.0002, Student’s unpaired t-test. n = 5 for controls and n = 7 for MCT2KO. At 6 weeks, (**), p =0.0040, Student’s unpaired t-test. n = 3 for controls and n = 5 for KO. **I-N**. Co-immunolabelling for GFP in green (I,L) and ACSS2 in red (J,M). K overlay of I-J. N overlay of M-N. I-J control. L-N MCT2 KO. White arrows indicate GFP+ ACSS2+ cells and are more numerous in controls. **O**. Quantification of the percentage of GFP+ cells expressing ACSS2 shows a significant decrease in MCT2 KO at 3 weeks, (*) p=0.0440, Student unpaired t-test and also at 6 weeks (**) p=0.0070, Student unpaired t-test. n=5 and 3 in controls at 3 and 6 weeks respectively and n=4 and N=5 in MCT2KO at 3 and 6 weeks respectively. Scale bars = 20 µm.

We then investigated the expression of another enzyme involved in lipid synthesis, acyl-coenzyme A synthetase short-chain family member 2 (ACSS2), which catalyzes the production of acetyl-CoA from acetate. Besides lipid synthesis, ACSS2 is crucial for energy generation and histone acetylation. We observed expression of ACSS2 oligodendrocytes in the Olig001-AAV transduced white matter (Fig.8I-K), some of which were GFP+ cells. In control mice, 64.52±8.23% of GFP+ cells were ACSS2+. However, in the MCT2KO mice, the proportion of ACSS2 expressing cells within the GFP population decreased to 26.73±13.96% at 3 weeks (Fig.8L-N,O). At 6 weeks, 53.33±11.59% of control GFP+ cells in the WM were ACSS2+, while in the MCT2KO these values decreased to 14.93±3.32% (Fig.8O). Therefore, our results show that demyelination after MCT2 deletion in oligodendrocytes is not due to oligodendroglial death but appears correlated to significantly reduced levels of two important enzymes that regulate the process of lipogenesis (FASN and ACSS2), suggesting a failure in myelin maintenance.

### Upregulation of LDHA in *MCT2KO* mice

We hypothesized that absence of MCT2 would reduce the ability of oligodendrocytes to import monocarboxylates and use them as metabolic fuels, forcing these cells to increase glucose metabolism. Thus, we then investigated the expression of lactate dehydrogenase A (LDHA), enzyme that converts pyruvate into lactate, the expression of which increases when glycolysis is upregulated. Gene expression data (*21*) and a recent study (*45*) show that oligodendrocytes under control conditions express very low levels of *ldha*. We performed co-labellings for LDHA and GFP in *control* vs *MCT2KO* mice at 6 weeks post Olig001-AAV injection. Both in *controls* and *MCT2KO* LDHA expression was observed on blood vessels (BV) and cells of macrophage/microglial morphology (Fig.9A-B). However, in *MCT2KO* but not in the *controls*, LDHA expression was also observed on a minor subset of GFP+ oligodendrocytes (Fig.9Bi). This expression was strong in a low proportion of cells (arrow in Fig.9Bi), and weaker in others (Fig.9Bi arrowheads). Quantification of the percentage of GFP-positive cells that were strongly positive for LDHA showed a significant difference between the control group and the MCT2KO group (*p = 0.0333), although this percentage still remained low (Fig.9C). Interestingly, the most striking difference in LDHA expression between the controls and MCT2KO was that, in the latter, strong LDHA expression was detected on axons positive for SMI32+, particularly the swollen ones (Fig.9Bii). Quantification revealed a significant increase in LDHA+ axons in MCT2KO mice (Fig.9D; (*p = 0.0199).

**Figure 9.**
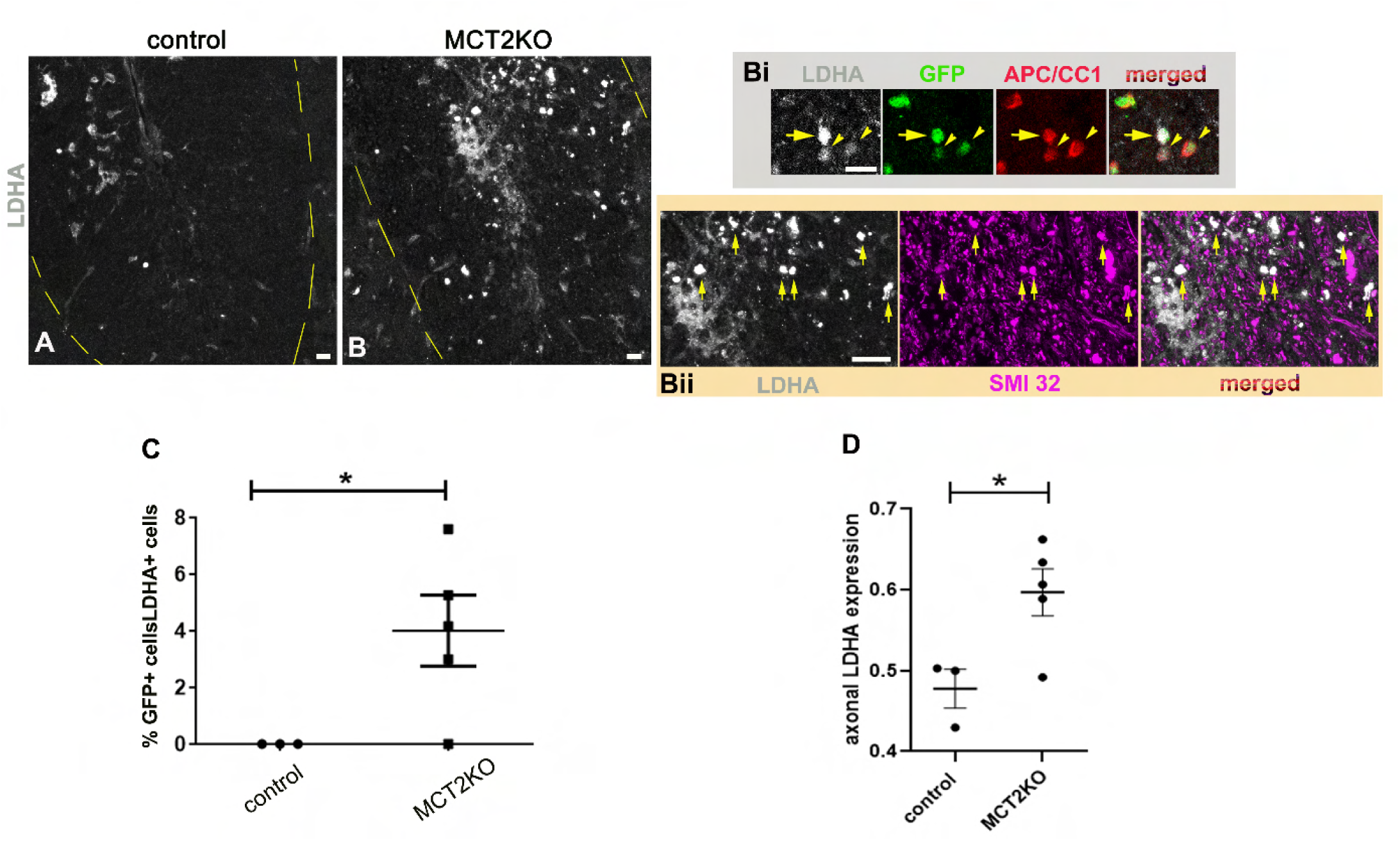
Increase in LDHA expression in MCT2KO. **A-B**. Immunolabelling for LDHA in in the dorsal funiculus of control (A) and MCT2KO (B) mice. Dashed lines delimit the white matter. **Bi**. Co-labelling for LDHA (gray), GFP (green) and APC/CC1 (red) reveals LDHA+, AAV-transduced oligodendrocytes in MCT2KO. Arrow indicates an MCT2KO oligodendrocyte strongly labelled by LDHA antibody, arrowheads indicate faintly labelled ones. **Bii**. Co-labelling for LDHA (gray) and SMI32 (magenta) reveals strong LDHA expression by SMI32+ (damaged) axons. Arrows indicate swollen SMI32+ axons that express LDHA. **C**. quantification of GFP+ cells strongly positive for LDHA reveals absence of colocalization in controls and an increase in MCT2KO mice (*p = 0.0333; unpaired t test with Welch’s correction). **D**. Significant increase in LDHA+ axons in MCT2KO mice (p= 0.0199, unpaired t test with Welch’s correction). n=3 for controls and n=5 for MCT2KO. Scale bars A, B, Bi 10µm, Bii 20µm.

We wondered whether LDHA upregulation in axons occurs because of demyelination, or because of metabolic alterations induced by oligodendroglial MCT2 deficiency. To investigate whether axonal LDHA is simply a consequence of demyelination, we investigated LDHA expression in LPC model of demyelination in the same area (dorsal funiculus). Within the demyelinated area at 14 days post lesion (dpl), we observed LDHA upregulation, with predominantly cellular staining pattern (Fig.S8A-B). Co-immunohistochemistry experiments showed these cells were macrophages (not shown). We did not observe axonal LDHA staining in the lesions. However, axonal pattern of staining was observed in the non-demyelinated tissue neighbouring the lesion (Fig.S8Bi-Biii). These observations suggests that axonal LDHA upregulation is not a simple consequence of demyelination.

Thus, our data indicate upregulation of LDHA in MCT2KO oligodendrocytes and in neighbouring axons, which is associated with axonal damage but does not appear to be a mere consequence of demyelination.

### Ketogenic diet attenuates pathological changes in *MCT2KO* white matter

We observed that MCT2 deletion in oligodendrocytes results in axonal damage associated with increased LDHA expression, the latter suggesting increased axonal glycolysis. Because increased neuronal glycolysis leads to neuronal dysfunction(*46*), we aimed to investigate whether exposure of mice to ketogenic diet, a low glucose diet, could alleviate axonal damage upon MCT2KO in oligodendrocytes, by reducing glucose availability and shifting neuronal metabolism towards ketone consumption, which was shown as neuroprotective (reviewed in(*15*)). Moreover, we also aimed to investigate whether MCT2KO oligodendrocytes can survive under low glucose conditions, when cells rely on alternative energy fuels such as monocarboxylates. We first quantified GFP cells to investigate the persistence of MCT2KO oligodendrocytes in under ketogenic diet. We did not observe significant differences between the two groups (GFP+ cells/mm^2^: 296±73.32 controls vs 436±113.7 MCT2KO, Fig.10A,D,G). Surprisingly, myelination by MCT2KO oligodendrocytes under ketogenic diet did not appear significantly affected (%MOG labelled area: 51.87±5.13 controls vs 50.4±3.24 MCT2KO; Fig.10B,E,H). LDHA labelling was scarce in both groups, detected only on a few cells with mononuclear morphology (Fig.S9). Importantly, quantification of SMI32 labelling revealed no significant differences between the 2 groups (% SMI32 labelled area 5.9±1.04 controls vs 5.69±2.95 MCT2KO, Fig.10I-K), and no significant differences were detected when quantifying CD45 (inflammatory cell marker) labelling either (% area labelled with CD45: 14.29±0.61 control vs 19.38±4.36 MCT2KO). Thus, ketogenic diet diminishes pathological features observed upon MCT2 deletion in oligodendrocytes.

**Figure 10.**
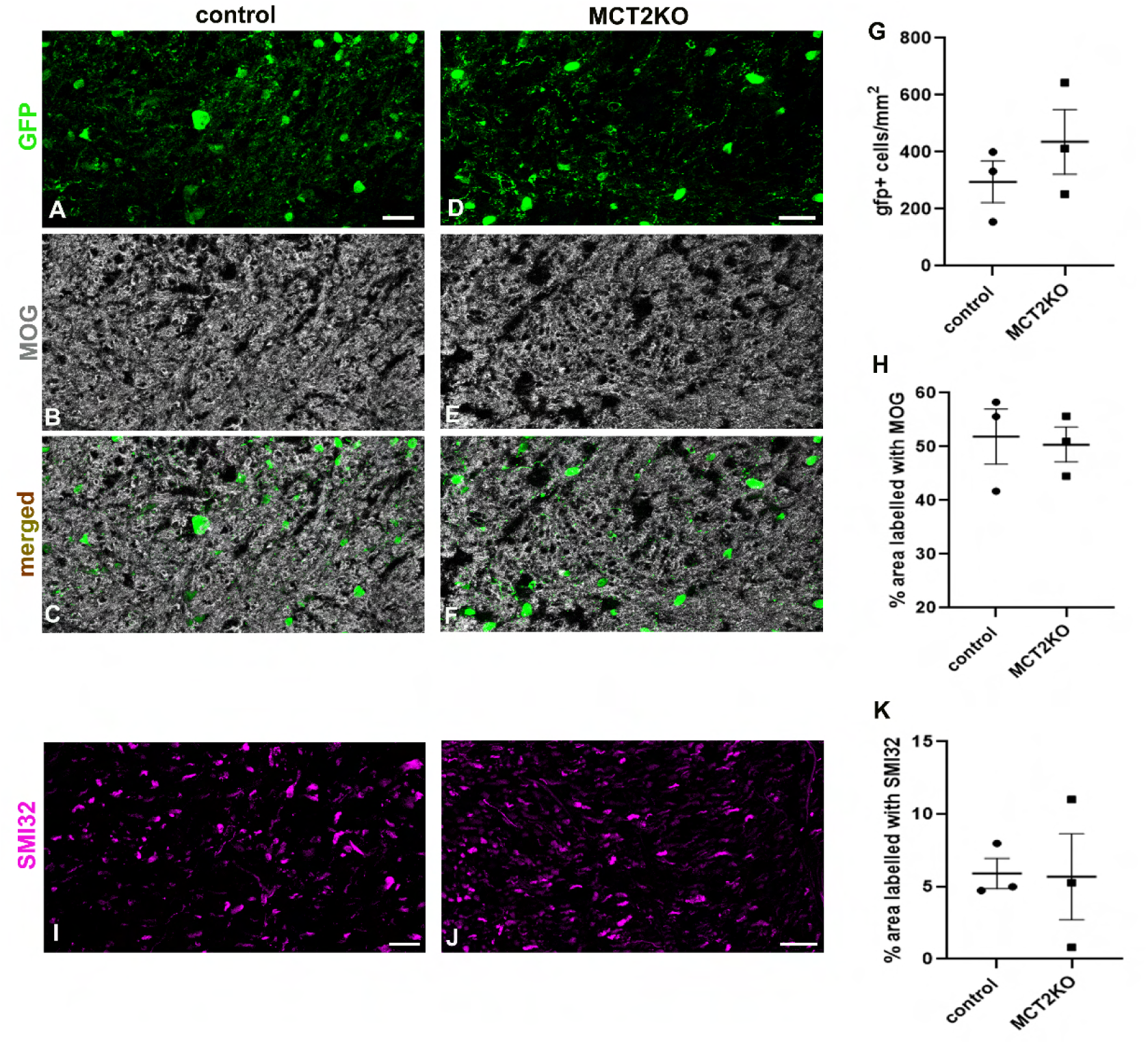
Effect of ketogenic diet on MCT2KO oligodendrocyte survival, demyelination and axonal damage. **A-F**. Co-immunohistochemistry for GFP (A,D) and myelin marker MOG (B,E) in control (A-C) and MCT2KO (D-F) mice. C-overlay of A-B, F-overlay of D-F. **G**. Quantification of GFP+ cells per area shows no significant differences between the groups. **H**. Quantification of MOG labelled area shows no significant differences between the groups. **I-J**. Labelling for SMI32 (non-phosphorylated neurofilament), a marker of axonal damage in control (I) and MCT2KO mice (J). **K**. Quantification of area labelled with SMI32 shows no differences between the groups. In all cases n=3 mice per group. Scale bars 20µm.

## DISCUSSION

The importance of MCT2 in neuronal function has been largely demonstrated. Thus, neuronal MCT2 expressed at the synapses imports astrocyte-derived lactate during the process of astrocyte-neuron lactate shuttle, and lactate is then metabolized to generate ATP that sustains synaptic transmission(*18*). Blocking MCT2 by single stranded oligonucleotide injection in the hippocampus(*47*) or deleting it specifically in hippocampal neurons(*38*) leads to neuronal dysfunction and impairments in memory. It is possible that MCT2 allows neurons to import not only lactate but also ketone bodies produced by astrocytic fatty acid oxidation, also shown as necessary to sustain neuronal function(*48, 49*). Thus, MCT2 is a crucial determinant of neuronal function by allowing these cells to import monocarboxylates with high affinity to sustain their metabolic needs.

Although oligodendrocytic expression of MCT2-encoding gene *Slc16a7* has been detected in mice(*19–21*) and humans(*22, 23*), potential function of MCT2 in myelinating oligodendrocytes has not been previously addressed. The expression of this high affinity monocarboxylate transporter by oligodendrocytes could empower these cells to efficiently import monocarboxylates under physiological conditions to sustain their extensive metabolic needs. Indeed, although synaptic activity is considered as the process that consumes most ATP in the brain(*9*), myelin synthesis also represents a significant energetic investment for the CNS(*10*). Besides large amounts of ATP needed to sustain the synthesis of lipids and proteins of the myelin sheath, oligodendroglial cells require enormous amounts of carbon precursors for the synthesis of myelin lipids (70-80% of myelin dry weight). Although during this process oligodendrocytes rely on astrocyte-derived lipids to an extent(*50*), myelin generation requires oligodendroglial fatty acid synthesis(*51*). It is important to highlight that active lipid synthesis is important not only during myelin generation but also during myelin maintenance, as interfering with lipid synthesis in adult myelinating oligodendrocytes disrupts myelin structure(*52*). Interestingly, literature highlights a significant contribution of both ketone bodies and lactate to myelin lipid synthesis, as ketone bodies are preferentially incorporated into myelin lipids over glucose during the development(*27*), and oligodendrocytes use lactate for lipid synthesis in vitro more than neurons or astrocytes(*26*). Whether monocarboxylate family contributes to lipid synthesis during myelin maintenance in vivo by mature oligodendrocytes is not clear, but our present data suggest this may be the case.

Myelinating oligodendrocytes have been considered for a long time as lactate producers rather than consumers, and as cells that predominantly rely on glucose consumption, the rate of which depends on axonal activity(*53, 54*). Thus, oligodendroglial glycolysis(*55*) and transfer of lactate via MCT1 to the axons (*56*) have been evoked as the basis of myelin maintenance and support of axonal integrity in the CNS. However, more recent reports suggest that oligodendroglial MCT1 is dispensable for these processes in non-aged mice(*30*). According to our present work, lack of myelination deficits in non-aged MCT1 oligodendroglial mutants could be explained by MCT2-mediated monocarboxylate import. Importantly, it has been recently shown that oligodendrocytes have very low expression of lactate dehydrogenases A and B, enzymes required to produce and consume lactate as metabolic fuels, respectively, as well as other glycolytic enzymes(*29*). Instead, it has been proposed that oligodendrocytes simply shuttle either the products of astrocytic glycolysis directly to the axons, and/or pyruvate produced by their own glycolysis(*29*). Here, we provide evidence that oligodendroglial metabolism and function might also require direct import of monocarboxylate family members via MCT2, such as lactate, pyruvate, ketone bodies, and acetate, all of which have been indicated as important metabolic supporters of the brain(*14, 16, 57, 58*). Thus, it could be that oligodendrocytes efficiently capture lactate via MCT2 from the extracellular space and shuttle it directly to the axons (as oligodendrocytes do not express LDH thus cannot metabolize lactate), which also may be the case for ketone bodies. Moreover, because oligodendrocytes express ketolytic enzymes(*19, 59*), MCT2 may also be used to efficiently capture ketone bodies (imported via endothelium from the blood, or produced and exported by astrocytes) as metabolic fuels for lipid and/or ATP synthesis (Fig.11A). Here we show that oligodendrocytes express ACSS2, an enzyme that converts acetate (another MCT2 substrate) to acetyl Co-A, and that both this enzyme and FASN (that catalyzes the first step of fatty acid synthesis) are downregulated after MCT2 deletion. Thus, our data directly support a link between MCT2, ACSS2 expression and fatty acid synthesis, although other mechanisms of MCT2 action mentioned above cannot be excluded.

**Figure 11.**
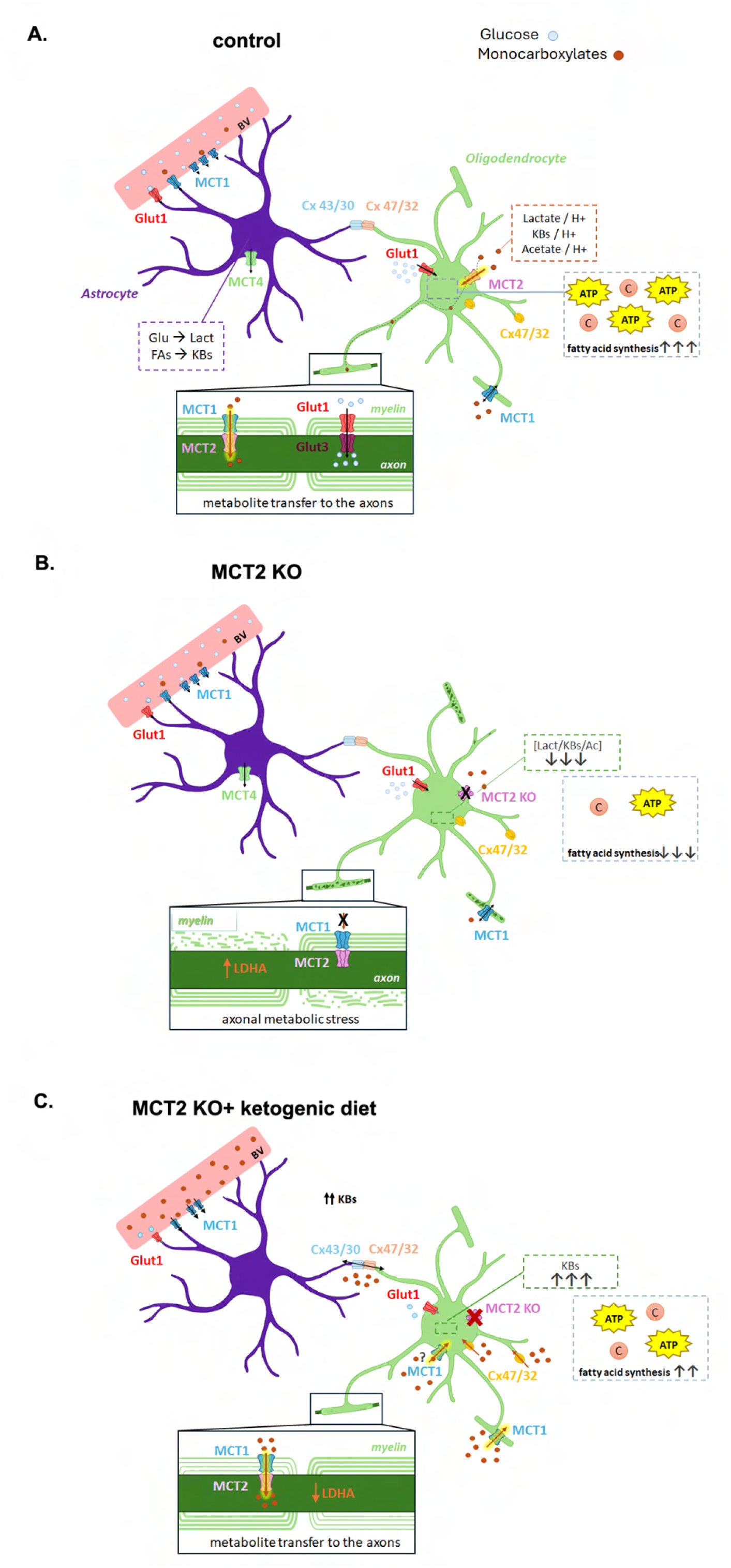
Schematic summary of myelinating oligodendrocyte metabolism and the role of MCT2. **A.** In the control white matter, glucose (Glu; light blue circles) and monocarboxylates (ketone bodies (KBs), lactate (Lact), acetate (Ac), pyruvate; brown circles) are delivered into the parenchyma via transporters expressed by endothelial cells in the blood vessels (BV). Glucose is imported by astrocytes and metabolized into pyruvate, which is then converted to lactate and shuttled outside astrocytes via MCT4(*18*). Astrocytes also metabolize fatty acids (FAs) to ketone bodies and export them via MCT4(*48*). Lactate(*100*) and ketone bodies can also be transferred to oligodendrocytes directly via connexin channels. MCT2 expressed by oligodendrocytes, given its high affinity, efficiently imports monocarboxylates from the extracellular space. Among these, ketone bodies, pyruvate and acetate can be metabolized to ATP or used as a carbon source for lipid synthesis. Lactate imported by oligodendrocytes is likely not metabolized to pyruvate because of low LDH expression(*29*) but may be shuttled via myelinic channels and MCT1 expressed on myelin to the axons that import it via MCT2. Oligodendrocytes also import glucose via Glut1, connexin hemichannels, or receive it from astrocytes via gap junctions (connexin channels)(*100*). Glucose then can be directly shuttled to the axons or metabolized to pyruvate, which may then be shuttled to the axons(*29*). **B**. MCT2 deletion in oligodendrocytes significantly diminishes the ability of oligodendrocytes to import monocarboxylates under standard conditions. This decreases both the availability of carbon molecules and ATP for lipid synthesis leading to metabolic deficit and failure of myelin maintenance. In addition, monocarboxylate delivery to the axon diminishes, leading to axonal LDHA upregulation and damage. **C.** Ketogenic diet increases the entry of ketone bodies to the CNS via MCT1 on endothelial cells. Moreover, it induces metabolic reprogramming of astrocytes allowing them to import ketone bodies via MCT1(*62*), which could then be delivered to oligodendrocytes via connexin channels. High extracellular ketone body concentrations also allow lower affinity transporters such as MCT1 or connexin hemichannels, to import these molecules into oligodendrocytes and sustain lipid synthesis and axonal support. Moreover, ketones can be directly imported and metabolized by neurons/axons(*62*).

Demyelination observed after MCT2 deletion in myelinating oligodendrocytes was not associated with oligodendrocyte death. These results resemble previous work showing that altered nutritional supply in cats and monkeys leads to myelin loss despite oligodendrocyte preservation(*60, 61*). Restoration of normal metabolite supply within a limited time frame in these models resulted in remyelination by pre-existing oligodendrocytes(*60, 61*). While we do not know whether re-expression of MCT2 by oligodendrocytes in our model would restore their myelinating capacity, we did observe that failure of myelin maintenance is attenuated if MCT2KO mice are subjected to a ketogenic diet. In this scenario, high ketone concentrations might allow for compensation of defective MCT2 by other MCTs or other transporters (e.g. connexin hemichannels). These transporters have lower affinity for monocarboxylates than MCT2 and thus likely cannot compensate for MCT2 deficiency under physiological conditions (low extracellular MC concentrations)(Fig.11B) but might be able to do so when extracellular ketone concentrations increase and those of glucose decrease, which is expected under ketogenic diet. Moreover, under ketogenic diet, astrocytes undergo metabolic reprogramming and import ketones(*62*), which could then be shuttled through connexin channels to oligodendrocytes (Fig.11C).

Besides myelination, another crucial function of oligodendrocytes in the CNS is the support of axonal integrity, dependent not only on the presence of myelin but also on its composition(*63–65*), presumably because molecular alterations in myelin can affect metabolite and organelle provision to the axons by oligodendrocytes via myelinic channels(*66–68*). We have observed that upon MCT2 deletion in oligodendrocytes, axons upregulate LDHA and SMI32, a marker of axonal damage. Although basal levels of neuronal LDHA expression appear required for fast axonal vesicular transport(*69*), LDHA upregulation by neurons has been previously correlated to increased angiogenesis and neurodegeneration(*70*), altered cognition in aged mice(*71*), as well as to elevated metabolic demand from chronic neuronal activation(*72*). LDHA upregulation likely reflects increased axonal glycolysis, that possibly takes place to compensate for deficient myelination and axonal support by oligodendrocytes that lack MCT2. Increased neuronal glycolysis leads to oxidative stress, as it redirects glucose utilization from pentose phosphate pathway (anti-oxidant) to pyruvate/lactate production, which results in neuronal dysfunction(*46*). Thus, it is possible that MCT2-deficient oligodendrocytes enter a “metabolite-sparing” state to sustain their own survival which decreases myelin lipid synthesis as well as metabolic provision to the axons, leading to increased axonal glycolysis and axonal damage, observed both immunohistochemically (increased SMI32 immunoreactivity) and by ultrastructural analyses.

Interestingly, we observed that LPC-induced demyelination (associated with oligodendrocyte death) does not lead to LDHA upregulation in demyelinated axons, suggesting that myelin loss itself does not upregulate axonal glycolysis. Instead, axonal LDHA was observed perilesionally. It is possible that presence of a hypermetabolic area (demyelinating lesion with hypercellularity and ongoing myelin repair) induces a certain degree of metabolic deprivation in the neighboring non-demyelinated white matter leading to aberrant axonal metabolism, perhaps because metabolite-deprived oligodendrocytes have impaired capacity to sustain axonal metabolic needs. On the other hand, within lesions where axons are demyelinated, these can import metabolites directly through specific transporters on their surface. This scenario is compatible with previous reports showing that myelin is a risk factor for the axons within inflamed environment(*73*), possibly because inflammatory cells are hypermetabolic(*11, 74*) and therefore may deplete myelinating oligodendrocytes thus indirectly myelinated axons, from metabolic fuels. Therefore, one could hypothesize that persistence of metabolically-altered oligodendrocytes (and sometimes myelin) is worse for the axons than oligodendrocyte death and short-term demyelination, provided it is followed by remyelination. Importantly, presence of dysfunctional oligodendrocytes could impair remyelination by OPCs as mature oligodendrocytes express netrin-1, which prevents OPC recruitment (*75–77*).

Importantly, axonal LDHA expression and damage were alleviated by ketogenic diet. Low glucose and direct neuronal import of ketone bodies during ketogenic diet likely decrease axonal glycolysis and enable neurons to increase their ketone metabolism, alleviating axonal damage. In line with our data showing axonal protection by ketogenic diet, exposure of mice with experimental autoimmune encephalomyelitis (EAE) to ketogenic diet was found to increase ketolytic metabolism in neurons and astrocytes and was neuroprotective(*62*).

It is important to highlight that monocarboxylates imported by MCT2 may fulfil functions other than that of ATP and/or lipid precursors. Lactate and ketone bodies have been shown as substrates for protein lactylation(*78*) and β-hydroxybutyrylation(*79*), respectively. Such modifications on histones can affect the expression of genes that regulate metabolism(*80*), which strongly suggests that even if oligodendrocytes do not express enzymes that allow them to convert lactate into pyruvate (that can be further metabolized in different metabolic pathways), lactate imported via MCT2 could still play an important role in regulating oligodendrocyte metabolism by executing epigenetic modifications. Such roles of lactate in oligodendrocytes should be addressed by future studies. Moreover, acetate and ketone body deprivation in MCT2-deficient oligodendrocytes could lead to a decrease in acetyl CoA, thus deficient histone (and other protein) acetylation, further affecting oligodendroglial function.

Our data show downregulation of *Slc16a7*/MCT2 expression by oligodendrocytes in progressive MS. We first performed *in silico* analyses of *Slc16a7* in different oligodendrocyte clusters described by Trobisch and colleagues(*22*), in control vs MS subjects. These data show predominant *Slc16a7* expression in the homeostatic cluster associated with white matter (SLC5A11 cluster). Cluster-specific enrichment for *Slc16a7* expression in human oligodendrocytes has also been reported by Sadick, ÒDea et al(*23*) who showed that *Slc16a7* expression is significantly enriched in oligodendrocyte cluster 2, the one associated with increased cholesterol (lipid) synthesis (https://liddelowlab.shinyapps.io/Sadick/). In addition, the analyses by Seeker et al.(*35*) showed *Slc16a7* expression in OligoC cluster, one of the 2 clusters enriched in the spinal cord (https://seeker-science.shinyapps.io/shiny_app_multi/). Interestingly, while our *in silico* analyses of Trobisch et al data(*22*) show that *Slc16a7* expression decreases in the white matter cluster in MS, this expression in cluster 2 was unchanged in Alzheimer’s disease(*23*). We then analyzed MCT2 protein expression in cerebellar tissue of control subjects vs patients with progressive MS using SOX10 as oligodendroglial marker. We analyzed three areas in MS tissue: NAWM, lesion border, and lesion core. Firstly, the number of SOX10+ cells at lesion borders varied between the patients in that some patients showed an increase at lesion borders as compared to the NAWM, while others showed a decrease. This may be due to the fact that SOX10 labels both oligodendrocytes and OPCs. While in the control WM and NAWM SOX10+ cells are predominantly mature oligodendrocytes and the minority are OPCs, the increase in SOX10+ cell numbers seen at some lesion borders may represent a block in OPC recruitment (reviewed in(*75*)). Importantly, all patients showed SOX10+ cell depletion in the lesion core, which is consistent with the idea that chronic demyelination in MS is associated with a strong decrease and sometimes even depletion of both OPCs and oligodendrocytes(*75*). Interestingly, the dynamics of oligodendroglial MCT2 expression did not follow that of oligodendroglial (SOX10+ cell) numbers. In the NAWM where large numbers of SOX10+ cells were detected, we observed a decrease in the percentage of these cells that express MCT2 compared to control subject white matter. In addition, the percentage of SOX10+ cells positive for MCT2 further decreased at lesion borders, suggesting that the proximity of lesions leads to MCT2 downregulation in oligodendroglia in progressive MS. This will likely impair the capacity of these cells to import monocarboxylates, which interferes with their capacity to synthesize lipids and sustain axonal metabolism, as suggested by our loss-of -function data in the mouse model. Thus, our data support the notion that oligodendroglial dysfunction in progressive MS is present beyond lesion areas leading to diffuse myelin/axonal damage in the NAWM that underlies “silent” disease progression(*81*).

Understanding MCT2 downregulation in MS would first require revealing the mechanisms that regulate physiological expression of this transporter in oligodendrocytes. In neurons, MCT2 expression appears regulated mainly at the translational level and involves molecules such as IGF-1/insulin(*82*), BDNF(*83, 84*) and noradrenaline(*85*). While it is not known whether these molecules also regulate oligodendroglial MCT2 expression, BDNF and noradrenaline have been reported to decrease in MS(*86, 87*). While transcriptional regulation of MCT2 was not previously reported in neurons, we have observed decreased oligodendrocyte *Slc16a7* mRNA expression in MS. Transcriptional regulation of MCT2 has been reported in cancer setting, as MCT2 overexpression observed in prostate cancer cells was observed to result from demethylation of the *Slc16a7* promoter, thus suggesting that epigenetic mechanisms regulate both mRNA and protein levels of MCT2(*88*). Decreased mRNA levels of *Slc16a7* were also reported in adipose tissue upon hypoxia(*89*). Interestingly, hypoxia was also associated with decreased MCT2 protein expression in brain tumors(*90*). Further studies should imperatively address the mechanisms underlying regulation of oligodendroglial MCT2.

Anomalies in brain metabolism are well recognized in MS and thought to play an important role in disease progression, which is why ensuring adequate metabolic support to stimulate myelin maintenance and repair might be crucial(*2, 11*). However, provision of adequate energy fuels will be effective only if corresponding transporters are expressed. Our work suggests that MCT2 is required for myelin maintenance and axonal support, but is downregulated on oligodendrocytes in MS, which suggests that these cells have an impaired capacity to perform a high affinity import of monocarboxylates, at least under physiological conditions. These findings therefore question the usefulness of potential monocarboxylate provision strategies (e.g. ketogenic diet, direct monocarboxylate supplementation etc.) as a supporting therapy in MS. However, in our mouse model, demyelination and axonal damage due to MCT2 deletion in oligodendrocytes were attenuated by ketogenic diet, likely due to direct axonal effects but also the fact that in situations of increased monocarboxylate (ketone) extracellular concentrations and low glucose, the high affinity transporter MCT2 is not indispensable because other transporters of lower affinity can capture and import these molecules. However, oligodendroglial and astrocytic connexins are also downregulated in chronic MS(*91*). Interestingly, our *in silico* analyses of *Slc16a1,* the gene encoding MCT1, a lower affinity MCT, show no changes in the white matter oligodendrocyte cluster in MS. Moreover, we observed increased expression of this gene in reactive oligodendrocyte clusters as compared to the homeostatic white matter cluster (the opposite of what was seen with *Slc16a7*). This suggests that, if MCT1 protein expression corresponds to that of mRNA, increased monocarboxylate supply may be effective in stimulating myelin preservation in MS as it would enable oligodendrocytes to import these molecules via MCT1. Thus, future studies should imperatively investigate potential changes in oligodendroglial MCT1 protein expression in MS NAWM to reveal whether metabolic therapies to increase monocarboxylate provision may be of interest to stimulate myelin maintenance in MS.

## MATERIALS AND METHODS

### Mice

Experiments with mice and rats were approved by the Animal Ethic Committees of the University of the Basque Country (UPV/EHU) and followed the European Communities Council Directive 2010/63/EU. Mice were housed in a laboratory environment with controlled temperature and 12 hours light/dark cycle, with food and water ad libitum. A previously described MCT2 floxed mouse line(*38*) and control C57Bl6J 4-6 months old mice were used for the experiments.

### AAV generation

The Cre-GFP construct(*92*) was purchased from Addgene (https://www.addgene.org/68544) and cloned into the oligodendrotropic AAV Olig001(*41*) generously provided by Sara Powell and Thomas McCown (University of North Carolina) at the University of North Carolina Vector Core.

### AAV injection

Spinal cord injections were performed as described previously(*76, 93*). Prior to the surgery mice were anaesthetized by an intraperitoneal injection of ketamine (90mg/kg) and xylazine (20mg/kg) cocktail in NaCl. Then, two longitudinal incisions into longissimus dorsi at each side of the vertebral column were performed to remove the tissue covering the column was removed. Animals were placed in a stereotaxic frame, the 13th thoracic vertebra was fixed in between the bars designed for manipulations on mouse spinal cord (Stoelting, Wood Dale, IL), and intervertebral space was exposed by removing the connective tissue. An incision into dura mater was performed using a 30-gauge needle, and 0.5 µL of AAV solution was injected using a glass micropipette attached via a connector to a Hamilton’s syringe and mounted on stereotaxic micromanipulator. The injection site was marked with sterile charcoal. The muscle sheaths were sutured with 3/0 Monocryl, and the skin incision was closed with sterile surgical clips.

MCT2^lox/lox^ mice injected with Olig001-Cre-GFP AAV are referred to as *MCT2KO*. Wildtype (*wt*) mice injected with Olig001-Cre-GFP and MCT2^lox/lox^ mice injected with Olig001-GFP were both used as controls. After confirming that Olig001-Cre-GFP injection in *wt* mice did not lead to pathological changes observed in *MCT2KO,* MCT2^lox/lox^ mice injected with Olig001-GFP were used as specific controls for quantification experiments, as those mice were from the same litters as *MCT2KO* and housed in the same ages, for we considered these as the best possible controls.

### Ketogenic diet experiments

Teklad custom ketogenic and control for KETO diets were purchased from Envigo. Ten days after AAV injection (minimum time required for AAV expression in the tissue), mice injected with Olig-001 AAV were switched to ketogenic diet (10% protein, <1% carbohydrate and 89% fat). Ketone levels in the blood were monitored using the GlucoMen Day METER 2K Blood Glucose/β-Ketone Monitoring System and corresponding ketone strips. Mice were sacrificed at 3 weeks post AAV injection (11 days after the initiation of the ketogenic diet).

### Perfusion and tissue processing

Mice were euthanized with an overdose of pentobarbital and perfused with a 2% paraformaldehyde (PFA; Electron Microscopy Sciences, Hatfield, PA) solution in phosphate-buffered saline (PBS, pH 7.4) for immunohistochemistry (IHC) experiments or a 2% PFA and 0.5% glutaraldehyde solution in 0.1M phosphate buffer (PB) for transmission electron microscopy (EM) experiments.

For IHC analyses, the brains and spinal cords were extracted and post-fixed for 20 minutes in 2% PFA. After rinsing 3 times with NaCl, the tissue was equilibrated in 15% sucrose, embedded in a 7% gelatine (type A porcine skin, Sigma)/15% sucrose solution in PBS, snap-frozen using isopentane (Sigma-Aldrich) at -55°C, and subsequently stored at -80°C prior to cryostat sectioning. Spinal cord cryosections, of 12 µm in thickness, were cut coronally on Super Frost Plus slides (Thermofisher) using the Leica CM1950 Cryostat. Slides were then air-dried for a minimum of 30 min and stored at -20°C.

For EM analyses, spinal cords were post-fixed overnight in 2% PFA and 0.5% glutaraldehyde, washed 3 times in 0.1M PB and stored in 0.1M PB + 0.05% Sodium azide prior to resin processing and embedding.

### Human Material

Human brain tissue was obtained from the UK and French MS Tissue Banks (Dr R. Reynolds, Imperial College, London and NeuroCEB, Pitié Salpêtrière, Paris respectively) and collected with the donors’ fully consent, following ethical approval.

Snap-frozen sections from postmortem cerebellum samples were analyzed to study the expression pattern of MCT2 and FASN. For MCT2, 4 controls and 4 MS patients were analyzed and for FASN there were 3 controls and 4 MS patients. The clinical features of these human subjects and lesions analyzed are detailed in Table S1.

### Immunohistochemistry (IHC)

Slides with spinal cord sections were air-dried at room temperature for 45min-1h and then rehydrated in TBS 1x for 30 min. Antigen retrieval was performed by heating the sections in a citrate-based retrieval buffer (Vector Laboratories/Dako) for 45 sec in a microwave. After serial washes, blocking buffer (0.1% Triton X-100, 10% goat serum in 1x TBS) was applied for 45 min, followed by incubation with primary antibodies (Table S2) diluted in the blocking buffer, overnight at 4°C. The next day, samples were washed and incubated with the secondary antibodies (Alexa-conjugated secondary antibodies, Thermofisher, used at 1:500) for an hour. Finally, slides were counterstained with DAPI, washed and mounted with Fluoromount-G.

For human tissue, histological assessment of the lesions was performed using Luxol Fast blue/Cresyl violet and Oil-red-O staining. Lesions were classified according to their inflammatory activity (KP1 and MHC-II immunolabelling) and on the basis of previously described histological criteria(*94, 95*) .

Cryostat sections (12 µm thickness) of snap-frozen control and MS cerebellum were rehydrated in PBS. Thereafter, slides were post-fixed for 20 minutes with PFA 2% and washed, to posteriorly microwave them in low-pH unmasking solution (Vector Laboratories). Sections were then preincubated in blocking buffer (10% normal goat serum, 0.1% Triton-X 100 in PBS) for 1 hour and incubated overnight with primary antibodies (Table S2) at 4°C. After overnight incubation, slides were extensively washed in PBS 0.1% Triton X-100 and incubated with appropriate secondary antibodies. For FASN immunohistochemistry, the signal was amplified using biotinylated secondary antibody followed by Alexa 488-conjugated streptavidin (Invitrogen). Nuclei were counterstained with Hoechst and slides mounted with Fluoromount-G.

### Mouse tissue imaging and quantification

Imaging of mouse tissue sections was performed using a Leica TCS STED CW SP8X confocal microscope and the Zeiss LSM 880 confocal microscope equipped with a 40x oil-immersion objective.

The area containing Olig-001 AAV-transduced cells was localized based on the expression of GFP reporter. Images were acquired at 40x and/or 63x magnification to cover the entire dorsal funiculus. At least two spinal cord levels separated by 144 µm minimum were imaged per mouse. Analyses were carried out using the ImageJ/Fiji software (NIH). Cells positive for GFP, for the marker of interest, and cells co-expressing GFP and the marker of interest were counted. The results are presented either as cells/mm^2^ or as the percentage of cells within the GFP+ population.

In the case of GFAP, MOG, MBP, and CD45 staining, a positivity threshold was set in the corresponding channel, and quantifications were performed as the percentage of area labelled within the ROI of interest. For colocalization studies, cells labeled on both channels were counted and the results were expressed as numbers of co-labelled cells per area or the percentage of the total.

In the case of neurofilaments, positivity thresholds were applied to sections labelled with anti-SMI31 (phosphorylated NF) or -SMI32 (non-phosphorylated NF) antibodies. The area labelled for these antigens was then quantified using the “area fraction” setting in Fiji. Numbers of labelled axons were quantified using “Analyze Particles” function in Fiji with the circularity set as 0.10-1.00, and lower particle area threshold of 1.5µm^2^ for regular axons, and 13 µm^2^ for swollen axons.

The analyses of axonal LDHA expression were performed on cross-sectioned axons using specific Fiji macros(*96*), except where specified. First, LDHA and SMI32 channels were extracted from multichannel z-stacks by selecting the plane with the highest SMI32 mean grey value (MGV) plus/minus 2 z-steps. ROIs were manually drawn in order to exclude areas where axons are longitudinally organized (rare, as the absolute majority of the axons in dorsal funiculus are coronally cut in transverse spinal cord sections), thus selecting the regions with circular orthogonally cut axons. In order to avoid manual bias while applying axonal masks, we applied the “Moments method threshold” colocalization quantification via BIOP-Jacop plugin(*97*), and calculated Manders coefficient, that was considered as quantification of “axonal LDHA expression” in Fig. 9D.

To illustrate myelin-axon status in the injected white matter, a 3D reconstruction of the colocalization between SMI31-MBP and GFP was generated using the Software Leica Application Suite X (LAS X).

### Electron microscopy

For EM processing, spinal cord area surrounding Olig001-AAV injection was coronally cut into 200 µm thick sections using a Leica VT1000S vibratome (Leica Biosystems, Wetzlar, Germany).

For pre-embedding immunostaining for GFP and Olig2 spinal cords were cut then into 50 µm slices. For aldehyde inactivation, sections were incubated in 1% sodium borohydride in PB and washed. Then, they were permeabilized by cryoprotection in 25% sucrose in PB followed by freeze-thaw cycles in methylbutanol. For the immunogold labeling, sections were incubated in blocking solution I (0.3% BSAc (Aurion) in PB) for an hour and then, anti-GFP (1:200; Aves Labs) primary antibody diluted in blocking solution was added from 36 up to 60h at 4°C and in agitation. Samples were washed after and incubated in blocking solution II (0.5% BSAc and 0.1% fish gelatin (Aurion) in PB) for an hour before incubation with the secondary antibody conjugated to colloidal gold (Aurion, 1:50) for 24h. Afterwards, for the silver enhancement, sections were washed and then immersed in equal volumes of reagents A and B (Aurion) in a dark chamber, to amplify the signal. Lastly, gold toning was performed by gold incubation, 0.05% gold chloride for 10 minutes, and posterior incubation in 0.3% sodium thiosulfate for two times.

For the embedding process, sections were washed with 0.1M PB and post-fixed with osmium tetroxide. Samples underwent a sequence of increasing ethanol concentrations washes for dehydration before uranyl acetate incubation. Propylene oxide was used after as an intermediate between alcohol washes and the final embedding in araldite. Semithin (1.5 um) sections were cut for identification of the area of interest, and ultrathin (70-80 nm) sections were obtained and stained with lead citrate for analysis.

The samples were examined and images were acquired at 80 kV on a FEI Tecnai G2 Spirit microscope (FEI Company, Hillsboro, OR) equipped with a Xarosa digital camera (20 megapixel resolution) using Radius image acquisition software (EMSIS GmbH, Münster, Germany).

TEM images were analyzed using Image J, and myelinated and demyelinated axons were counted.

### Quantification of MCT2 and SOX10 in human samples

All tissues obtained from multiple sclerosis patients consisted of parts of the cerebellum, where different types of lesions were found. For the quantification of MCT2 and SOX10, images were acquired in the active zones of 3 active lesions, 2 chronically active lesions, and 1 shadow plaque lesion.

Images were acquired in the core region of each lesion, in the perilesion, and in the normal-appearing white matter. These areas were identified based on Luxol fast Blue, Cresyl violet, and MHCII staining. Images were acquired in these areas using a camera (AxioCam; Zeiss, Jena, Germany) attached to a Zeiss Cell Observer with Apotome module (Zeiss).

SOX10+ and MCT2+ cells, and co-labeled cells within the lesion core, were quantified as the number of positive cells per unit area or as percentage of the total SOX10+ cells.

### Statistical analyses

GraphPad Prism 8 software was used to perform all statistical analyses (GraphPad Software, CA, USA). Data were presented as the mean ± standard error of the mean (SEM). All data sets were tested for normality and homoscedasticity. For the comparison of two groups, if data passed normality test, parametric unpaired t-test with Welch’s correction was applied; if data did not pass the normality test, Mann-Whitney’s test was applied. Comparisons between more than two groups were performed according to the normality test results either using one-way Analysis of variance (ANOVA), or in the case of negative normality test, using Kruskal Wallis test. Statistical significance was represented as p<0.05 (*), p<0.01 (**) and p<0.001 (***).

### In silico analyses of Slc16a7 and Slc16a1 expression

Single nucleus transcriptomic count matrix and metadata information were retrieved from https://ms-cross-regional.cells.ucsc.edu (accession date: 20 November 2023; downloaded object: integrated seurat.rds), corresponding to the publication of Trobisch et al. 2022(*22*). Bioinformatic analyses were conducted with the R programming language (version 4.1.2) (https://www.R-project.org). First, processing was performed following the standard Seurat R pipeline (package version 5.0.1.)(*98*). We selected nuclei annotated as oligodendrocytes that met the quality control criteria established by the original authors. We retained nuclei that expressed at least 250 different genes, had a minimum of 400 counts, and contained less than 5% mitochondrial genes. Normalization was performed with the NormalizeData function, and gene expression levels were visualized using the VlnPlot function. The differential expression analysis for *Slc16a7* gene across groups was performed with DESeq2(*99*). The pseudobulk raw counts were generated by Seurat’s AggregateExpression function, on which we applied the negative binomial generalized linear model, adjusting for age when indicated in the results section. For multiple comparison testing, p-values were adjusted by the Benjamini and Hochberg procedure (Benjamini and Hochberg, 1995), and significantly differential expression defined as false discovery rate < 0.05.

## Data availability

The data that support the findings of this study are available from the corresponding author upon reasonable request.

## ACKNOWLEDGEMENTS

We thank Dr. Laura Escobar (Achucarro Basque Center for Neuroscience), Dr. Ricardo Andrade (Microscopy Platform UPV/EHU), Susana González Granero (Cavanilles Institute/University of Valencia), Mario Soriano (Centro de Investigación Principe Felipe), and Luis Manuel Mendoza (Achucarro Basque Center for Neuroscience) for technical assistance.

We are grateful to Drs Sara Powell and Thomas McCown (Un. of North Carolina) for generously providing Olig001 plasmid.

This work was supported by: Ministerio de Ciencia, Innovación y Universidades-España (SAF2015-74332-JIN and PID2022-143020OB-I00 to VT, and PID2023-152688OB-I00 to CM); ARSEP Foundation and France Sclèrose en Plaques Foundation to VT and BNO, Consellería de Educación, Universidades y Empleo-Generalitat Valenciana (CIDEXG/2023/23) to VT, BIOEF-EITB-Maratoia (BIO23/EM/008) to VT, NeurATRIS to VT and BNO, CIBERNED (CB06/05/0076 to CM and CB06/05/1131 to JMGV), and Gobierno Vasco (IT1203-19 and IT-1551-22 to CM). LIU was a recipient of the Basque Government PhD studentship. ISS is supported by a predoctoral grant FPU20/03544 funded by the Spanish Ministry of Universities. FGG is supported by PID2021-124430OA-I00 funded by MCIN/AEI/10.13039/501100011033 and by “ERDF A way of making Europe”, and by CIAICO/2023/149 funded by the Consellería de Educación, Cultura, Universidades y Empleo de la Generalitat Valenciana.

## REFERENCES

1. S. Camandola, M. P. Mattson, Brain metabolism in health, aging, and neurodegeneration. EMBO J 36, 1474–1492 (2017).

2. R. M. Heidker, M. R. Emerson, S. M. LeVine, Metabolic pathways as possible therapeutic targets for progressive multiple sclerosis. Neural Regen Res 12, 1262–1267 (2017).

3. A. M. Amorini et al., Serum lactate as a novel potential biomarker in multiple sclerosis. Biochim Biophys Acta 1842, 1137–1143 (2014).

4. G. Lazzarino et al., Serum Compounds of Energy Metabolism Impairment Are Related to Disability, Disease Course and Neuroimaging in Multiple Sclerosis. Mol Neurobiol 54, 7520–7533 (2017).

5. M. E. Witte, D. J. Mahad, H. Lassmann, J. van Horssen, Mitochondrial dysfunction contributes to neurodegeneration in multiple sclerosis. Trends Mol Med 20, 179–187 (2014).

6. D. Jakimovski et al., Multiple sclerosis. Lancet 403, 183–202 (2024).

7. C. Lubetzki, B. Zalc, A. Williams, C. Stadelmann, B. Stankoff, Remyelination in multiple sclerosis: from basic science to clinical translation. Lancet Neurol 19, 678–688 (2020).

8. L. Sokoloff, in Handbook of Physiology, American Physiological Society, F. J., M. H. W., H. V. E, Eds. (American Physiological Society, Washington DC, 1960), vol. 3, pp. 1843–1864.

9. J. J. Harris, R. Jolivet, D. Attwell, Synaptic energy use and supply. Neuron 75, 762–777 (2012).

10. J. J. Harris, D. Attwell, The energetics of CNS white matter. J Neurosci 32, 356–371 (2012).

11. V. Tepavčević, Oligodendroglial Energy Metabolism and (re)Myelination. Life (Basel) 11, (2021).

12. M. Bradl, H. Lassmann, Oligodendrocytes: biology and pathology. Acta Neuropathol 119, 37–53 (2010).

13. K. Pierre, L. Pellerin, Monocarboxylate transporters in the central nervous system: distribution, regulation and function. J Neurochem 94, 1–14 (2005).

14. E. Benarroch, What Is the Role of Lactate in Brain Metabolism, Plasticity, and Neurodegeneration? Neurology 102, e209378 (2024).

15. H. Yang, W. Shan, F. Zhu, J. Wu, Q. Wang, Ketone Bodies in Neurological Diseases: Focus on Neuroprotection and Underlying Mechanisms. Front Neurol 10, 585 (2019).

16. Q. He et al., Acetate enables metabolic fitness and cognitive performance during sleep disruption. Cell Metab 36, 1998–2014.e1915 (2024).

17. L. Pellerin, L. H. Bergersen, A. P. Halestrap, K. Pierre, Cellular and subcellular distribution of monocarboxylate transporters in cultured brain cells and in the adult brain. J Neurosci Res 79, 55–64 (2005).

18. L. Pellerin, P. J. Magistretti, Sweet sixteen for ANLS. J Cereb Blood Flow Metab 32, 1152–1166 (2012).

19. A. Zeisel et al., Molecular Architecture of the Mouse Nervous System. Cell 174, 999–1014.e1022 (2018).

20. J. D. Cahoy et al., A transcriptome database for astrocytes, neurons, and oligodendrocytes: a new resource for understanding brain development and function. J Neurosci 28, 264–278 (2008).

21. Y. Zhang et al., An RNA-sequencing transcriptome and splicing database of glia, neurons, and vascular cells of the cerebral cortex. J Neurosci 34, 11929–11947 (2014).

22. T. Trobisch et al., Cross-regional homeostatic and reactive glial signatures in multiple sclerosis. Acta Neuropathol 144, 987–1003 (2022).

23. J. S. Sadick et al., Astrocytes and oligodendrocytes undergo subtype-specific transcriptional changes in Alzheimer’s disease. Neuron 110, 1788–1805.e1710 (2022).

24. G. Chakraborty, P. Mekala, D. Yahya, G. Wu, R. W. Ledeen, Intraneuronal N-acetylaspartate supplies acetyl groups for myelin lipid synthesis: evidence for myelin-associated aspartoacylase. J Neurochem 78, 736–745 (2001).

25. A. M. Krasnow, D. Attwell, NMDA Receptors: Power Switches for Oligodendrocytes. Neuron 91, 3–5 (2016).

26. L. I. Sánchez-Abarca, A. Tabernero, J. M. Medina, Oligodendrocytes use lactate as a source of energy and as a precursor of lipids. Glia 36, 321–329 (2001).

27. J. W. Koper, M. Lopes-Cardozo, L. M. Van Golde, Preferential utilization of ketone bodies for the synthesis of myelin cholesterol in vivo. Biochim Biophys Acta 666, 411–417 (1981).

28. J. E. Rinholm et al., Regulation of oligodendrocyte development and myelination by glucose and lactate. J Neurosci 31, 538–548 (2011).

29. E. Späte et al., Downregulated expression of lactate dehydrogenase in adult oligodendrocytes and its implication for the transfer of glycolysis products to axons. Glia 72, 1374–1391 (2024).

30. T. Philips et al., MCT1 Deletion in Oligodendrocyte Lineage Cells Causes Late-Onset Hypomyelination and Axonal Degeneration. Cell Rep 34, 108610 (2021).

31. P. K et al., Enhanced expression of three monocarboxylate transporter isoforms in the brain of obese mice. The Journal of physiology 583, (2007).

32. V. Governa et al., Landscape of surfaceome and endocytome in human glioma is divergent and depends on cellular spatial organization. Proc Natl Acad Sci U S A 119, (2022).

33. Y. Zhang et al., Purification and Characterization of Progenitor and Mature Human Astrocytes Reveals Transcriptional and Functional Differences with Mouse. Neuron 89, 37–53 (2016).

34. P. K, M. PJ, P. L, MCT2 is a major neuronal monocarboxylate transporter in the adult mouse brain. Journal of cerebral blood flow and metabolism : official journal of the International Society of Cerebral Blood Flow and Metabolism 22, (2002).

35. L. A. Seeker et al., Brain matters: unveiling the distinct contributions of region, age, and sex to glia diversity and CNS function. Acta Neuropathol Commun 11, 84 (2023).

36. Y. Osanai, R. Yamazaki, Y. Shinohara, N. Ohno, Heterogeneity and regulation of oligodendrocyte morphology. Front Cell Dev Biol 10, 1030486 (2022).

37. J. Edmond, Ketone bodies as precursors of sterols and fatty acids in the developing rat. J Biol Chem 249, 72–80 (1974).

38. C. Netzahualcoyotzi, L. Pellerin, Neuronal and astroglial monocarboxylate transporters play key but distinct roles in hippocampus-dependent learning and memory formation. Prog Neurobiol 194, 101888 (2020).

39. J. S. Francis et al., N-acetylaspartate supports the energetic demands of developmental myelination via oligodendroglial aspartoacylase. Neurobiol Dis 96, 323–334 (2016).

40. R. J. Mandel et al., Novel oligodendroglial alpha synuclein viral vector models of multiple system atrophy: studies in rodents and nonhuman primates. Acta Neuropathol Commun 5, 47 (2017).

41. S. K. Powell et al., Characterization of a novel adeno-associated viral vector with preferential oligodendrocyte tropism. Gene Ther 23, 807–814 (2016).

42. B. D. Trapp et al., Axonal transection in the lesions of multiple sclerosis. N Engl J Med 338, 278–285 (1998).

43. M. P. Pender, K. B. Nguyen, P. A. McCombe, J. F. Kerr, Apoptosis in the nervous system in experimental allergic encephalomyelitis. J Neurol Sci 104, 81–87 (1991).

44. M. P. Pender, M. J. Rist, Apoptosis of inflammatory cells in immune control of the nervous system: role of glia. Glia 36, 137–144 (2001).

45. S. E et al., Downregulated expression of lactate dehydrogenase in adult oligodendrocytes and its implication for the transfer of glycolysis products to axons. Glia, (2024).

46. D. Jimenez-Blasco et al., Weak neuronal glycolysis sustains cognition and organismal fitness. Nat Metab 6, 1253–1267 (2024).

47. A. Suzuki et al., Astrocyte-neuron lactate transport is required for long-term memory formation. Cell 144, 810–823 (2011).

48. B. Morant-Ferrando et al., Fatty acid oxidation organizes mitochondrial supercomplexes to sustain astrocytic ROS and cognition. Nat Metab 5, 1290–1302 (2023).

49. M. Guzmán, C. Blázquez, Is there an astrocyte-neuron ketone body shuttle? Trends Endocrinol Metab 12, 169–173 (2001).

50. N. Camargo et al., Oligodendroglial myelination requires astrocyte-derived lipids. PLoS Biol 15, e1002605 (2017).

51. P. Dimas et al., CNS myelination and remyelination depend on fatty acid synthesis by oligodendrocytes. Elife 8, (2019).

52. X. Zhou et al., Mature myelin maintenance requires Qki to coactivate PPARβ-RXRα-mediated lipid metabolism. J Clin Invest 130, 2220–2236 (2020).

53. A. S. Saab et al., Oligodendroglial NMDA Receptors Regulate Glucose Import and Axonal Energy Metabolism. Neuron 91, 119–132 (2016).

54. Z. J. Looser et al., Oligodendrocyte-axon metabolic coupling is mediated by extracellular K. Nat Neurosci 27, 433–448 (2024).

55. U. Fünfschilling et al., Glycolytic oligodendrocytes maintain myelin and long-term axonal integrity. Nature 485, 517–521 (2012).

56. Y. Lee et al., Oligodendroglia metabolically support axons and contribute to neurodegeneration. Nature 487, 443–448 (2012).

57. A. Tiwari et al., Mitochondrial pyruvate transport regulates presynaptic metabolism and neurotransmission. Sci Adv 10, eadp7423 (2024).

58. N. J. Jensen, H. Z. Wodschow, M. Nilsson, J. Rungby, Effects of Ketone Bodies on Brain Metabolism and Function in Neurodegenerative Diseases. Int J Mol Sci 21, (2020).

59. E. R. Saito et al., Alzheimer’s disease alters oligodendrocytic glycolytic and ketolytic gene expression. Alzheimers Dement 17, 1474–1486 (2021).

60. I. D. Duncan, A. Brower, Y. Kondo, J. F. Curlee, R. D. Schultz, Extensive remyelination of the CNS leads to functional recovery. Proc Natl Acad Sci U S A 106, 6832–6836 (2009).

61. I. D. Duncan et al., The adult oligodendrocyte can participate in remyelination. Proc Natl Acad Sci U S A 115, E11807–E11816 (2018).

62. T. Düking et al., Ketogenic diet uncovers differential metabolic plasticity of brain cells. Sci Adv 8, eabo7639 (2022).

63. J. M. Edgar et al., Oligodendroglial modulation of fast axonal transport in a mouse model of hereditary spastic paraplegia. J Cell Biol 166, 121–131 (2004).

64. J. M. Edgar et al., Early ultrastructural defects of axons and axon-glia junctions in mice lacking expression of Cnp1. Glia 57, 1815–1824 (2009).

65. C. Lappe-Siefke et al., Disruption of Cnp1 uncouples oligodendroglial functions in axonal support and myelination. Nat Genet 33, 366–374 (2003).

66. J. M. Edgar et al., Río-Hortega’s drawings revisited with fluorescent protein defines a cytoplasm-filled channel system of CNS myelin. J Anat 239, 1241–1255 (2021).

67. K. J. Chapple et al., A myelinic channel system for motor-driven organelle transport. bioRxiv, 2024.2006.2002.591488 (2024).

68. N. Snaidero et al., Antagonistic Functions of MBP and CNP Establish Cytosolic Channels in CNS Myelin. Cell Rep 18, 314–323 (2017).

69. M. Mc Cluskey, H. Dubouchaud, A. S. Nicot, F. Saudou, A vesicular Warburg effect: Aerobic glycolysis occurs on axonal vesicles for local NAD+ recycling and transport. Traffic 25, e12926 (2024).

70. H. Lin et al., Extracellular Lactate Dehydrogenase A Release From Damaged Neurons Drives Central Nervous System Angiogenesis. EBioMedicine 27, 71–85 (2018).

71. A. K. Frame et al., Altered neuronal lactate dehydrogenase A expression affects cognition in a sex- and age-dependent manner. iScience 27, 110342 (2024).

72. A. Ksendzovsky et al., Chronic neuronal activation leads to elevated lactate dehydrogenase A through the AMP-activated protein kinase/hypoxia-inducible factor-1α hypoxia pathway. Brain Commun 5, fcac298 (2023).

73. E. Schäffner et al., Myelin insulation as a risk factor for axonal degeneration in autoimmune demyelinating disease. Nat Neurosci 26, 1218–1228 (2023).

74. C. Schiepers et al., Positron emission tomography, magnetic resonance imaging and proton NMR spectroscopy of white matter in multiple sclerosis. Mult Scler 3, 8–17 (1997).

75. V. Tepavčević, C. Lubetzki, Oligodendrocyte progenitor cell recruitment and remyelination in multiple sclerosis: the more, the merrier? Brain, (2022).

76. V. Tepavčević et al., Early netrin-1 expression impairs central nervous system remyelination. Ann Neurol 76, 252–268 (2014).

77. J. M. Bin et al., Full-Length and Fragmented Netrin-1 in Multiple Sclerosis Plaques Are Inhibitors of Oligodendrocyte Precursor Cell Migration. Am J Pathol, (2013).

78. X. Liu, Y. Zhang, W. Li, X. Zhou, Lactylation, an emerging hallmark of metabolic reprogramming: Current progress and open challenges. Front Cell Dev Biol 10, 972020 (2022).

79. T. Zhou et al., Function and mechanism of histone β-hydroxybutyrylation in health and disease. Front Immunol 13, 981285 (2022).

80. F. Jing, J. Zhang, H. Zhang, T. Li, Unlocking the multifaceted molecular functions and diverse disease implications of lactylation. Biol Rev Camb Philos Soc, (2024).

81. I. V. Allen, S. McQuaid, M. Mirakhur, G. Nevin, Pathological abnormalities in the normal-appearing white matter in multiple sclerosis. Neurol Sci 22, 141–144 (2001).

82. J. Chenal, K. Pierre, L. Pellerin, Insulin and IGF-1 enhance the expression of the neuronal monocarboxylate transporter MCT2 by translational activation via stimulation of the phosphoinositide 3-kinase-Akt-mammalian target of rapamycin pathway. Eur J Neurosci 27, 53–65 (2008).

83. C. Robinet, L. Pellerin, Brain-derived neurotrophic factor enhances the expression of the monocarboxylate transporter 2 through translational activation in mouse cultured cortical neurons. J Cereb Blood Flow Metab 30, 286–298 (2010).

84. C. Robinet, L. Pellerin, Brain-derived neurotrophic factor enhances the hippocampal expression of key postsynaptic proteins in vivo including the monocarboxylate transporter MCT2. Neuroscience 192, 155–163 (2011).

85. K. Pierre, R. Debernardi, P. J. Magistretti, L. Pellerin, Noradrenaline enhances monocarboxylate transporter 2 expression in cultured mouse cortical neurons via a translational regulation. J Neurochem 86, 1468–1476 (2003).

86. N. Karimi et al., Blood levels of brain-derived neurotrophic factor (BDNF) in people with multiple sclerosis (MS): A systematic review and meta-analysis. Mult Scler Relat Disord 65, 103984 (2022).

87. A. Torrillas-de la Cal et al., Chemogenetic activation of locus coeruleus neurons ameliorates the severity of multiple sclerosis. J Neuroinflammation 20, 198 (2023).

88. N. Pertega-Gomes et al., Epigenetic and oncogenic regulation of SLC16A7 (MCT2) results in protein over-expression, impacting on signalling and cellular phenotypes in prostate cancer. Oncotarget 6, 21675–21684 (2015).

89. F. Pérez de Heredia, I. S. Wood, P. Trayhurn, Hypoxia stimulates lactate release and modulates monocarboxylate transporter (MCT1, MCT2, and MCT4) expression in human adipocytes. Pflugers Arch 459, 509–518 (2010).

90. C. Cheng et al., Alterations of monocarboxylate transporter densities during hypoxia in brain and breast tumour cells. Cell Oncol (Dordr) 35, 217–227 (2012).

91. C. Papaneophytou, E. Georgiou, K. A. Kleopa, The role of oligodendrocyte gap junctions in neuroinflammation. Channels (Austin) 13, 247–263 (2019).

92. P. J. Kennedy et al., Class I HDAC inhibition blocks cocaine-induced plasticity by targeted changes in histone methylation. Nat Neurosci 16, 434–440 (2013).

93. M. S. Aigrot et al., Genetically modified macrophages accelerate myelin repair. EMBO Mol Med, e14759 (2022).

94. C. F. Lucchinetti, W. Brück, M. Rodriguez, H. Lassmann, Distinct patterns of multiple sclerosis pathology indicates heterogeneity on pathogenesis. Brain Pathol 6, 259–274 (1996).

95. T. Kuhlmann et al., An updated histological classification system for multiple sclerosis lesions. Acta Neuropathol 133, 13–24 (2017).

96. J. Schindelin et al., Fiji: an open-source platform for biological-image analysis. Nat Methods 9, 676–682 (2012).

97. S. Bolte, F. P. Cordelières, A guided tour into subcellular colocalization analysis in light microscopy. J Microsc 224, 213–232 (2006).

98. T. Stuart et al., Comprehensive Integration of Single-Cell Data. Cell 177, 1888–1902.e1821 (2019).

99. M. I. Love, W. Huber, S. Anders, Moderated estimation of fold change and dispersion for RNA-seq data with DESeq2. Genome Biol 15, 550 (2014).

100. N. Meyer et al., Oligodendrocytes in the Mouse Corpus Callosum Maintain Axonal Function by Delivery of Glucose. Cell Rep 22, 2383–2394 (2018).

