## Supplemental material for "Monocarboxylate transporter 2 is required for the maintenance of myelin and axonal integrity by oligodendrocytes"

### SUPPLEMENTAL FIGURES

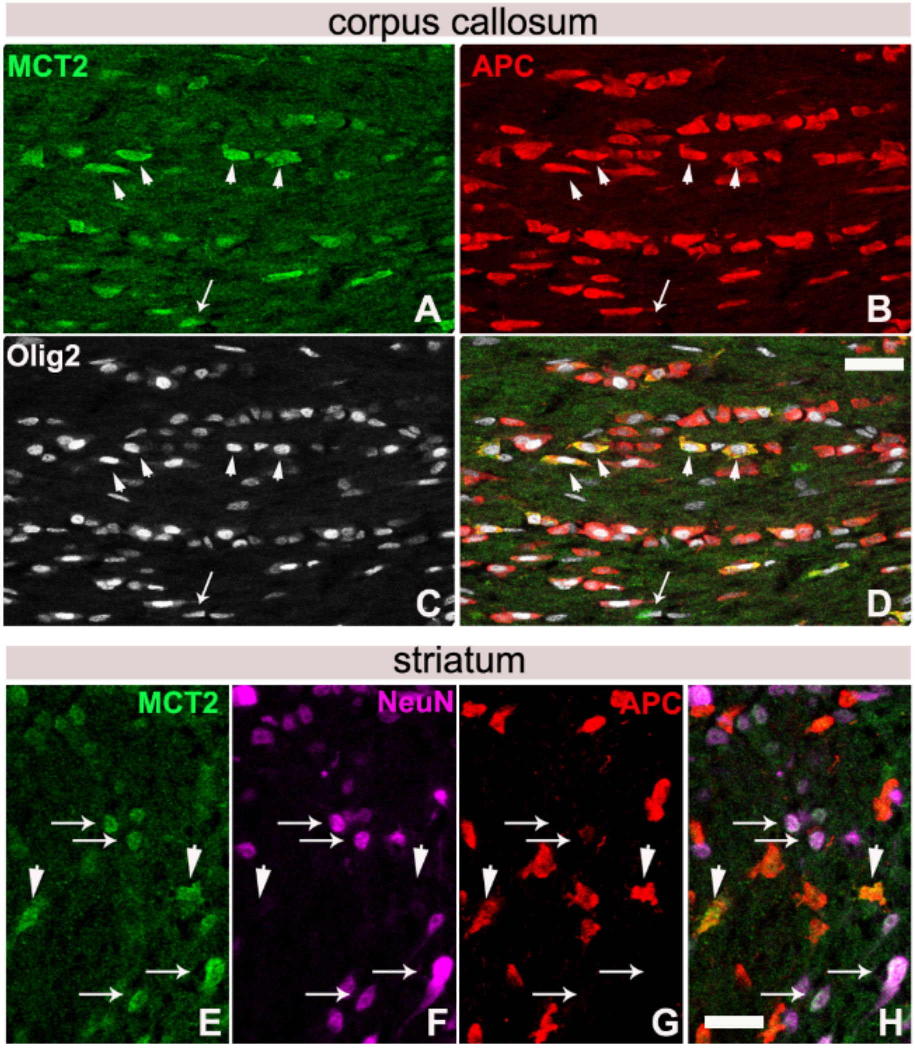


**Figure S1. Expression of MCT2 in the mouse brain**. Co-immunolabellings for MCT2 (green), APC/CC1 (red), Olig2 (white), and NeuN (purple). **A-D**. In the corpus callosum (white matter), MCT2 is seen on mature oligodendrocytes (APC+Olig2+, white arrowheads) and OPCs (Olig2+APC-,white arrows). **E-H**. In the striatum, MCT2+ cells are NeuN+ neurons (white arrows) and APC+ oligodendrocytes. Scale bar 20µm.

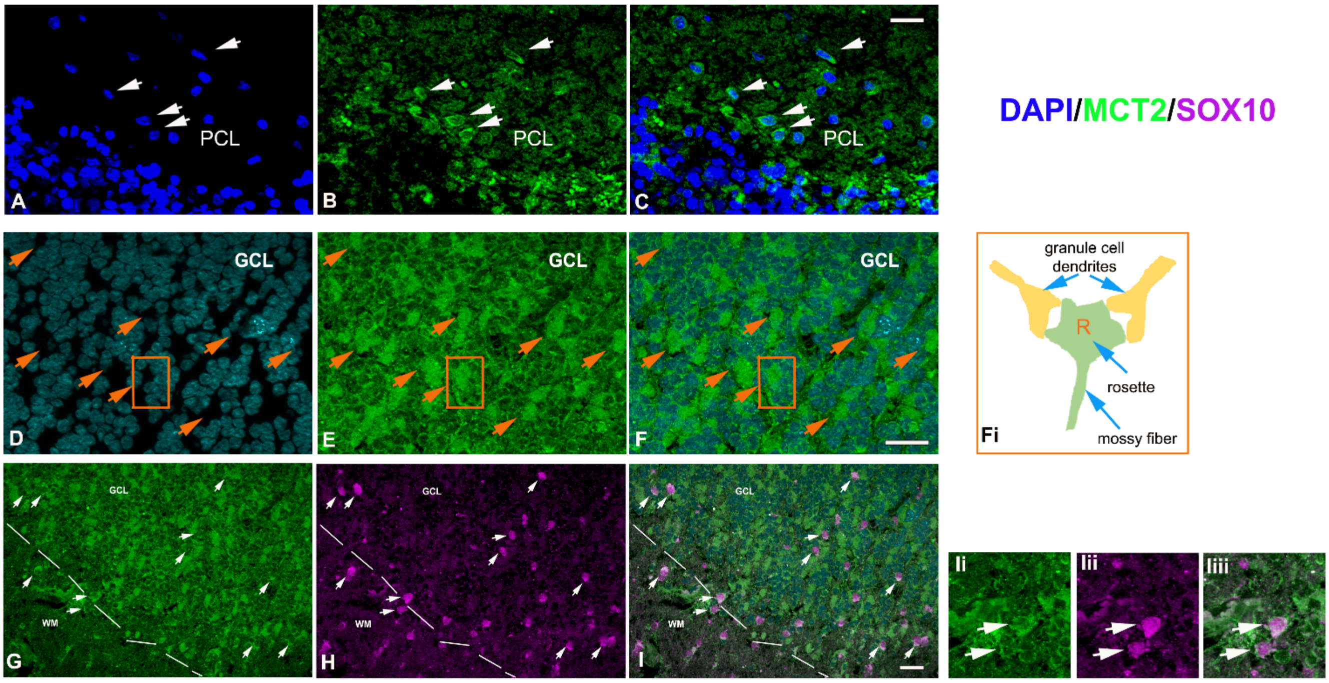


**Figure S2. Expression of MCT2 in the human cerebellum.** Blue-DAPI, green-MCT2, magenta-Sox10 (oligodendroglial marker). **A-C**. MCT2 positivity is observed on neurons in the Purkinje cell layer (PCL). **D-F**. In the granule cell layer (GCL), strong MCT2 positivity is observed in DAPI-free areas, corresponding to rosettes in the glomeruli (orange arrowheads), structures rich in synapses between granule cell dendrites and pre-synaptic terminals of mossy fibers (illustrating schema in Fi). **G-I**. MCT2 labelling is also observed on Sox10+ cells in the GCL, although stronger labelling is observed in the neighboring white matter (WM). Arrows indicate double-labelled cells. Scale bars 20 µm.

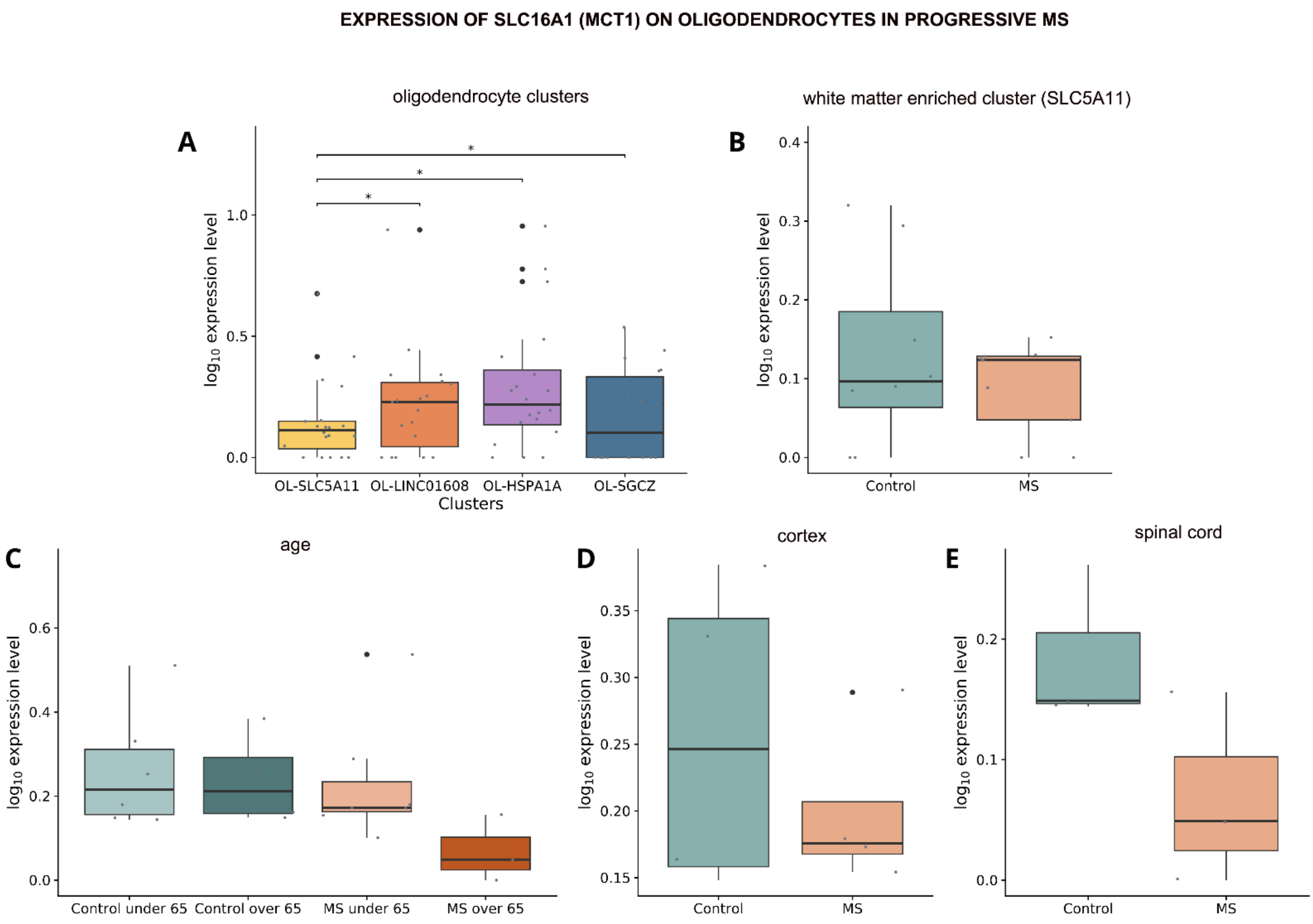


**Figure S3**. Analyses of *Slc16a1* expression changes in progressive MS. **A-E**. Normalized and logarithmically scaled *Slc16a1* gene expression values across different comparisons: **A.** oligodendrocyte clusters, **B**. SLC5A11 white matter-enriched oligodendrocyte cluster, **C**. stratification by age groups, **D.** cerebral cortex, and **E.** spinal cord. Boxplots represent the median (horizontal line) and interquartile range (box). Whiskers extend to the smallest and largest values within 1.5 times the interquartile range. Outliers are shown as large black dots, while small gray dots represent individual samples. MS: multiple sclerosis; OL: oligodendrocytes.

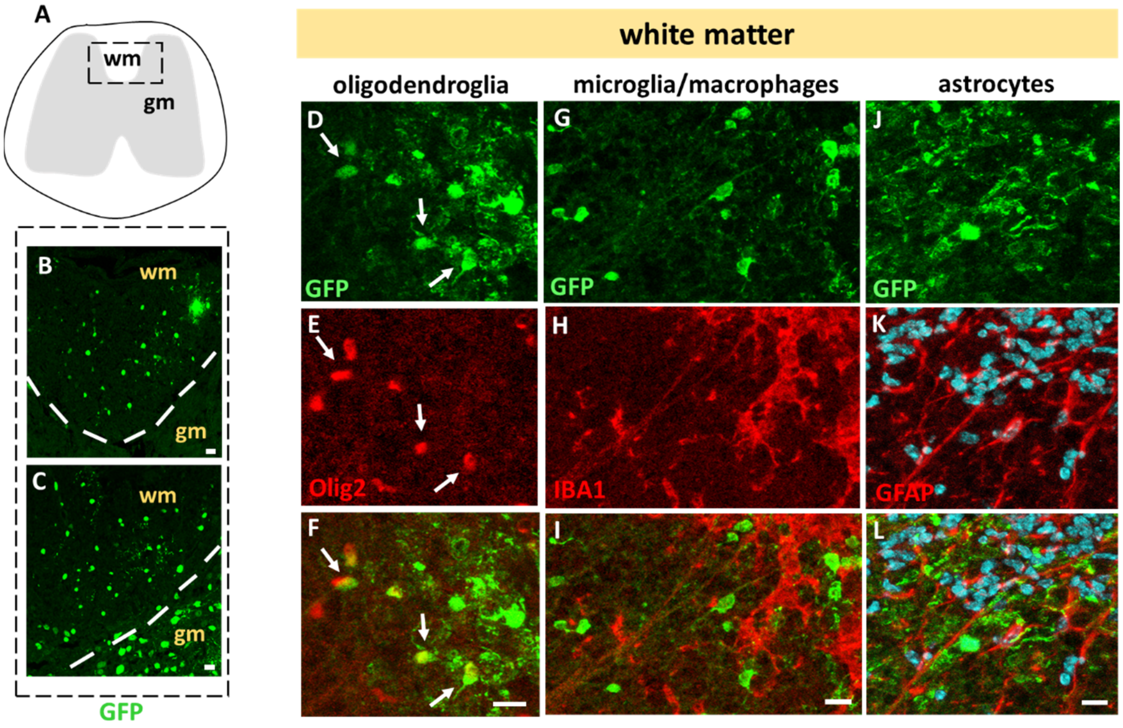


**Figure S4. Tropism of Olig001-AAV. A**. Schematic representation of the coronal section of the spinal cord. The rectangle indicates the area shown in B and C. **B-C**. Immunolabelling for GFP reveals Olig001-AAV transduced cells. **B**. In a subset of animals, these are found exclusively in the white matter (wm). **C**. In others, GFP staining is also present in the neighboring gray matter (gm). D-L. Characterization of GFP+ cells in Olig001-AAV-transduced white matter. **D-F**. Co-labelling for GFP in green (D) and oligodendroglial marker Olig2 in red (E). F. Overlay of D-E. Arrows indicate double-labelled cells. **G-I**. Co-labelling for GFP in green (G) and microglia/macrophage marker IBA1 in red (H). I. Overlay of G-H. **J-L**. Co-labelling for GFP in green (J) and astrocyte marker GFAP in red (K). L. Overlay of J-K. Scale bars 10 µm.

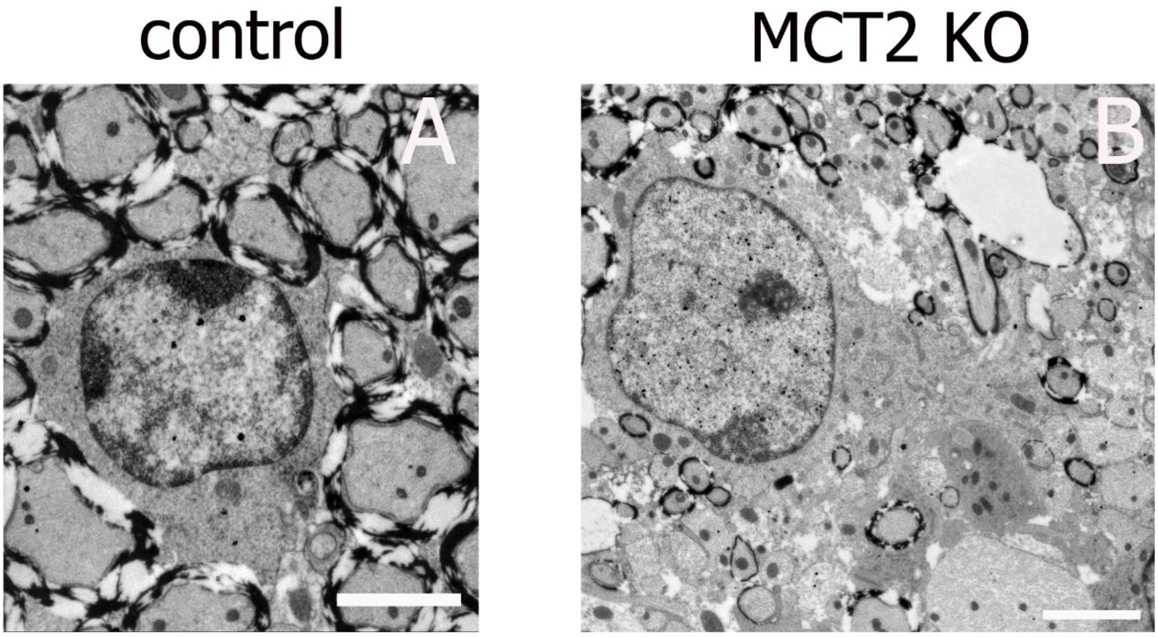


**Figure S5. Immunogold labelling for Olig2 in control versus MCT2 KO oligodendrocytes. A**. Olig2+ cell in control mouse. **B**. Olig2+ cell in MCT2 KO mouse. Scale bars 2µm.

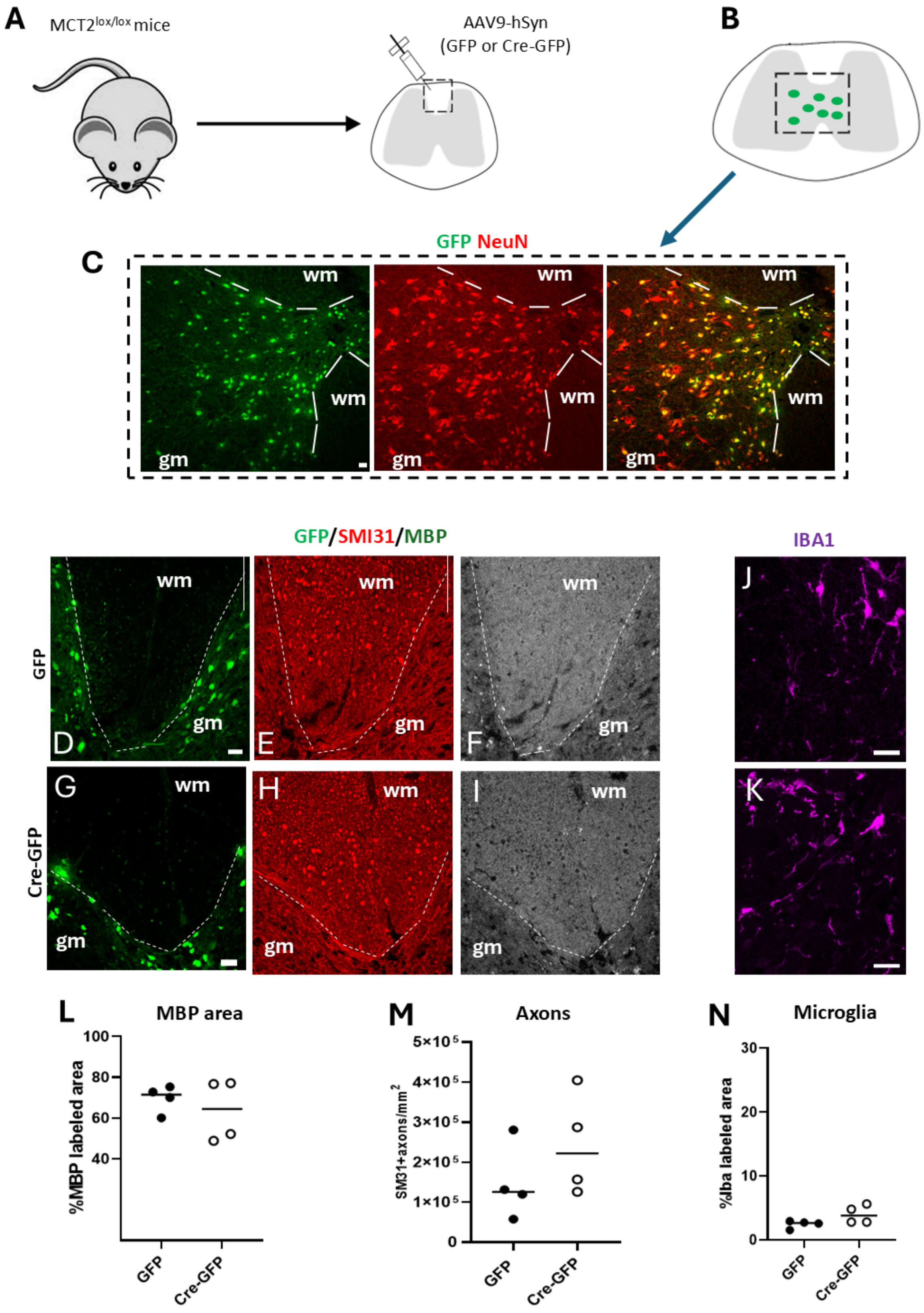


**Figure S6. MCT2 deletion in spinal cord neurons does not lead to demyelination in the neighboring white matter. A**. Schematic summary of the experimental approach employed to delete *Slc16a7* (MCT2) in spinal cord neurons: MCT$^{lox/lox}$ mice were injected with AAV-hSyn-GFP or AAV-hSyn-Cre-GFP in the dorsal white matter, using the same coordinates as those for Olig001-AAV injection. **B**. Schematic presentation of the coronal section of the spinal cord indicating the location of GFP+ cells exclusively in the gray matter. **C**. Co-labelling for GFP (green) and neuronal marker NeuN (red) shows that all AAV-transduced cells are neurons. **D-I**. Co-labelling for GFP in green (D,G), axonal marker SMI31 in red (E,H), and myelin marker MBP in gray (F,I) in the AAV-hSyn-GFP- injected (D-F) and AAV-hSyn-Cre-GFP-injected (G-I) $MCT^{lox/lox}$ mouse. **J-K**. Labelling for microglia/macrophage marker IBA1 in the AAV-hSyn-GFP- injected (J) and AAV-hSyn-Cre-GFP-injected (K) $MCT^{lox/lox}$ mouse. Quantitative comparison of the MBP-labelled area (**L**), SMI31+ axon number (**M**), and IBA1 labelled area (**N**) shows no differences between the groups. Scale bars 20 µm. gm-gray matter, wm-white matter.

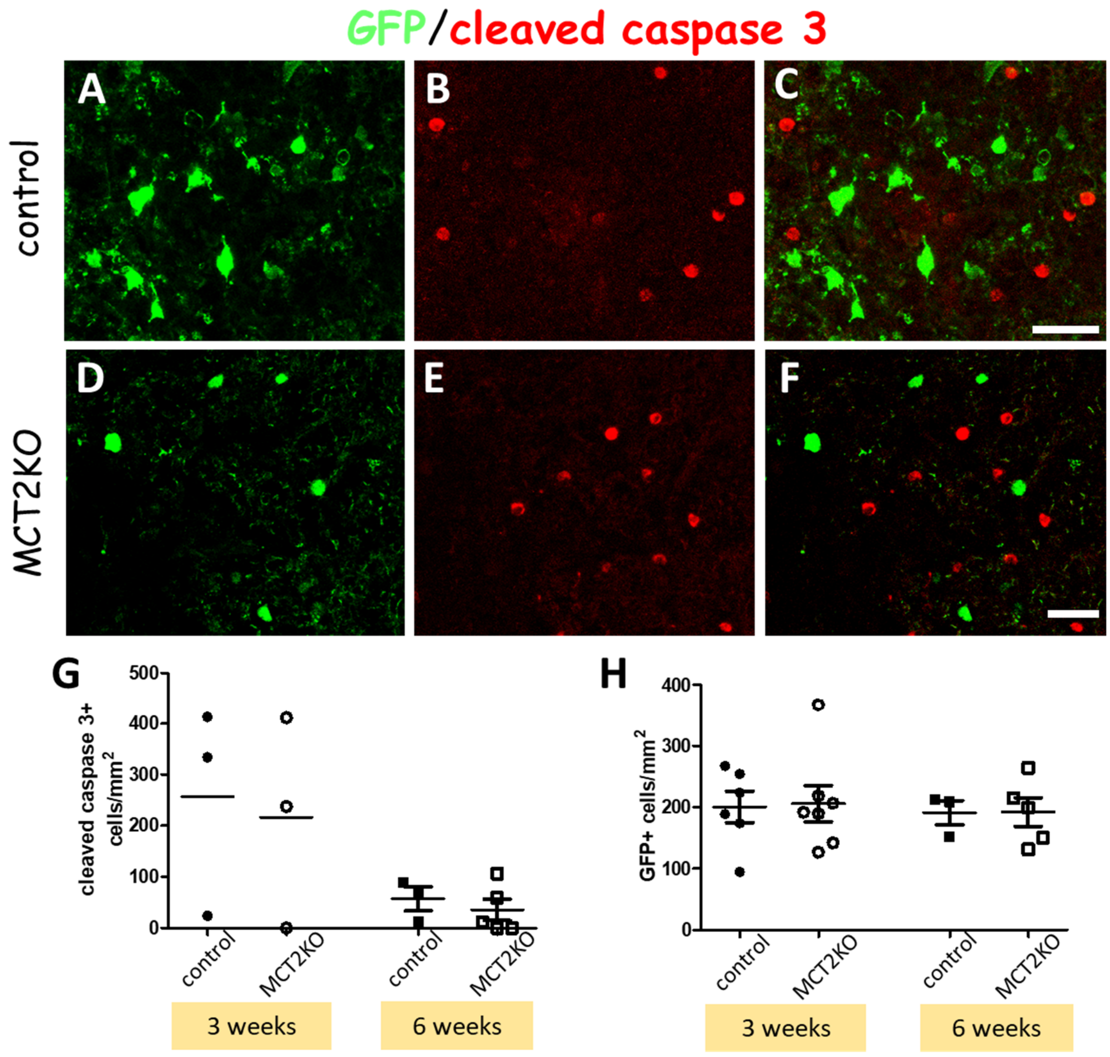


**Figure S7. MCT2 deletion does not lead to transduced cell death. A-F**: Sections of dorsal funiculus of $MCT2^{lox/lox}$ mice sacrificed at 3 weeks post Olig001-AAV injection co-labelled for GFP in green (A,D) and cleaved caspase 3 in red (B,E). C. Overlay of A-B. F. Overlay of D-E. A-C control mouse. D-F. MCT2KO mouse. Absence of colocalization of GFP and cleaved caspase 3 in both cases. **G**. Quantification for casp3+ cells per area shows no differences between the groups. n=3 for all conditions except for MCT2KO at 6 weeks where n=5. **H**. Quantification for GFP+ cells per area shows no differences between the groups. n=6 for controls at 3 weeks, n=7 for MCT2KO at 3 weeks, n=3 for controls at 6 weeks and n=5 for MCT2KO at 6 weeks. Scale bars=20µm.

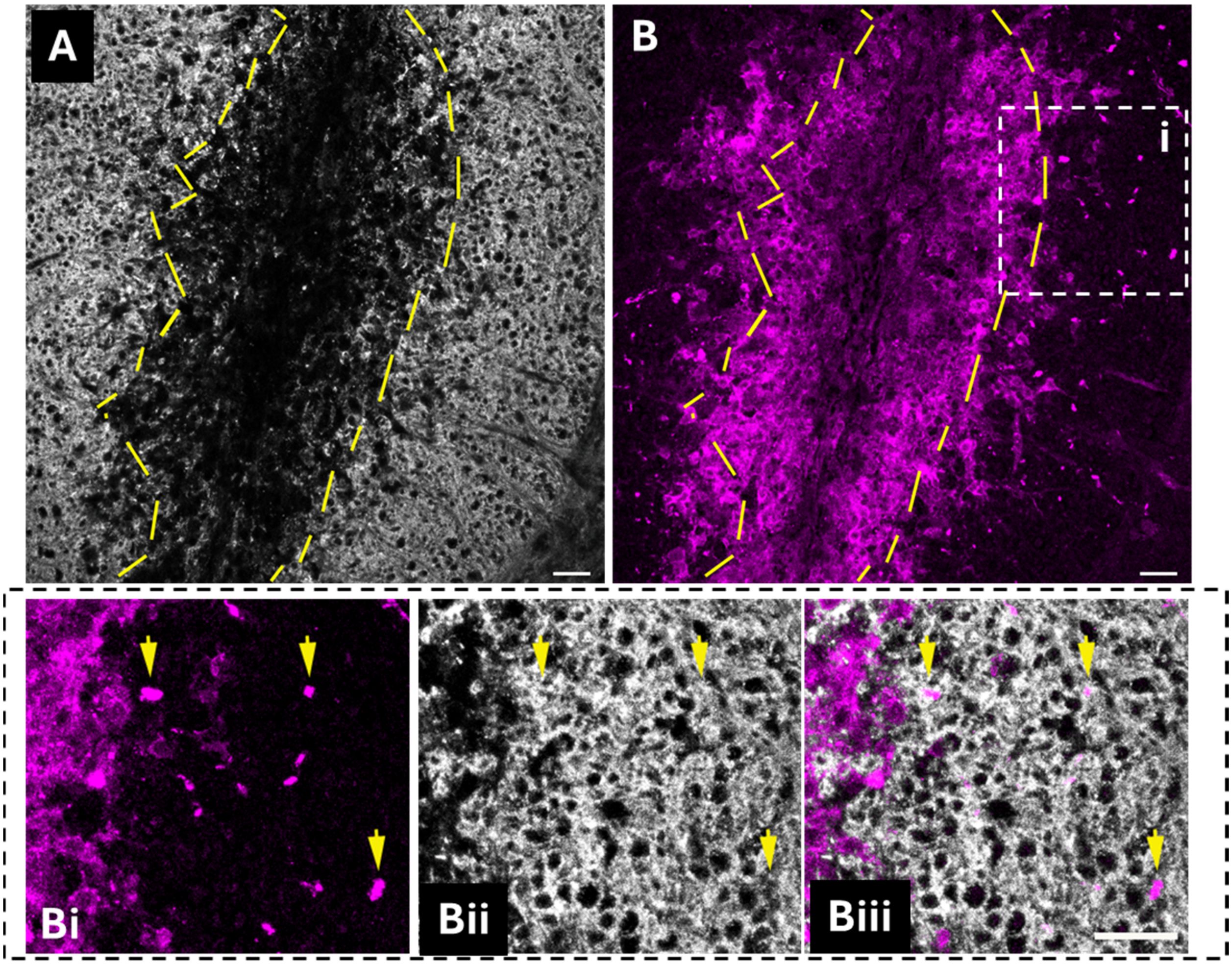


**Figure S8. LDHA is expressed by axons in perilesion after LPC induced demyelination**. **A-B** Images of demyelinating lesion induced by lysolecithin (LPC) injection in the wt mouse spinal cord, at 14 days post lesion (dpl). Co-immunolabelling for myelin marker MOG (white) and LDHA (magenta). Dashed lines delimit the area originally demyelinated. Some faint MOG staining is observed at the lesion border indicating ongoing remyelination. While LDHA labelling inside the demyelinated area shows cellular pattern, axonal staining pattern is observed only in the myelinated tissue adjacent to the lesion. **Bi-Biii** higher magnification of the area indicated by white rectangle in B. Arrows indicate LDHA labelled particles located within the myelin sheath (axons). Scale bars=20µm.

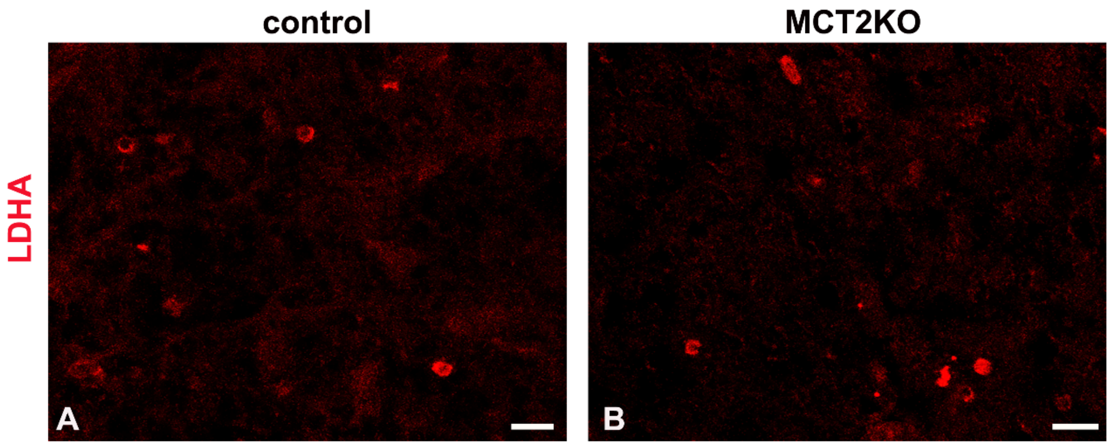


**Figure S9. LDHA staining in control and MCT2KO mice exposed to ketogenic diet. A**. Control mouse spinal cord tissue. **B**. MCT2KO mouse spinal cord tissue. In both, a few mononuclear-like cells labelled with LDHA are present, but axonal staining is not observed. Scale bar 10 µm.

### SUPPLEMENTAL TABLES

| Patients | | Sex | Age (at death) | Disease duration (y) | Disease course | Lesion activity |
|---|---|---|---|---|---|---|
| **MS 121** | | Female | 49 | 14 | PRMS | Chronic Active |
| **MS 74** | | Female | 64 | 36 | SPMS | Active |
| **MS 76** | MS76-CLC1 | Female | 49 | 18 | SPMS | Shadow Plaque/ Chronic Active |
| | MS76-CLB2 | | | | | Shadow Plaque |
| **MS 122** | | Male | 44 | 10 | SPMS | Active |
| **C 026** | | Female | 78 | Myeloid leukaemia | | |
| **C025** | | Male | 35 | Carcinoma of the tongue | | |
| **C015** | | Male | 82 | Chronic schizophrenia | | |
| **C022** | | Female | 69 | Lung cancer | | |

**Table S1. Clinical features of control subjects and patients with MS whose cerebellar tissue was analysed in the study.**

| Antibody | Host | Dilution | Reference |
|---|---|---|---|
| **Anti-MCT 2** | Mouse/Rabbit | 1:50/1:150 | Santa Cruz Biotech., sc-14924/Alomone labs., AMT-012 |
| **Anti-Olig 2** | Mouse IgG2a | 1:300 | Millipore, MABN50 |
| **Anti-APC-CC1** | Mouse IgG2b | 1:150 | Millipore OP80 |
| **Anti-NeuN** | Mouse | 1:500 | Sigma-Aldrich MAB377 |
| **Anti-GFP** | Chicken/Rabbit/Rat | 1:500 | Aves Labs., GFP-1020/Millipore AB3080/ Nacalai 04404-84 |
| **Anti-Cre** | Mouse | 1:250 | Millipore MAB3120 |
| **Anti-Iba 1** | Guinea Pig | 1:300 | Synaptic Systems 234 308 |
| **Anti-GFAP** | Mouse | 1:500 | Millipore MAB360 |
| **Anti-SMI 31** | Mouse IgG1 | 1:500 | Biolegend 801602 |
| **Anti-SMI 32** | Mouse | 1:500 | Biolegend 801701 |
| **Anti-MBP** | Chicken | 1:150 | Millipore ab9348 |
| **Anti-MOG** | Mouse IgG1 | 1:300 | Millipore MAB5680 |
| **Anti-Casp 3** | Rabbit | 1:400 | Cell Signaling Tech., 9664 |
| **Anti-FASN** | Rabbit | 1:200 | Abcam ab22759 |
| **Anti-ACSS2** | Rabbit | 1:300 | Abcam ab133664 |
| **Anti-LDH A** | Rabbit | 1:150 | Cloud-Clone Corp. PAB370Mu01 |
| **Anti-CD45** | Rat | 1:100 | Invitrogen 14-0451-82 |
| **Anti-CD3** | Rat IgG1 | 1:25-1:50 | Bio-Rad MCA500G |
| **Anti-MCT2 (For human tissue)** | Rabbit | 1:200 | Thermofisher-PA5-52225 |
| **Anti-SOX10** | Goat | 1:200 | R&D Systems AF2864 |
| **Anti-MOG (For human tissue)** | Mouse | 1:10 | Hybridoma supernatant (Glasgow,UK) |

**Table S2. Primary antibodies used for immunofluorescence.**
